# fastACCORD enables ultrahigh-dimensional partial correlation modeling for multi-omic data integration

**DOI:** 10.64898/2026.09.20.753046

**Authors:** Sungdong Lee, Qing Zhao, Dongwon Kim, Sang-Yun Oh, Joong-Ho Won, Hyungwon Choi

## Abstract

Gene co-expression networks reflect transcription factor-associated regulation together with epigenetic influences such as DNA methylation, chromatin states, and histone modifications. To distinguish gene-gene dependencies that persist after accounting for methylation covariation, large-scale statistical inference conditioning on hundreds of thousands of molecular features is necessary, but the task remains computationally intractable for conventional Gaussian graphical modelling approaches. Here we present fastACCORD, a scalable computational framework for ultrahigh-dimensional partial correlation modeling. fastACCORD combines row-separable optimization, ℓ2 stabilization, and a semismooth Newton solver in a PyTorch implementation for CPU and CUDA-enabled GPU hardware. We applied it to matched transcriptomic and methylomic profiles from 16 TCGA cancer types and obtained joint networks containing >300,000 molecular features per cancer. The resulting multi-omic networks revealed cancer-specific methylation-expression dependencies, including recurrent methylation-associated expression repression of metabolic genes. Methylation-adjusted gene co-expression networks were sparser than networks estimated from mRNA data alone, but more enriched for ChIP-seq-supported TF-target relationships and curated co-regulon annotations. TF-target subnetworks further revealed cancer-specific architectures consistent with lineage identity, oncofetal reactivation, and tumor microenvironment-associated programs. Together, these analyses establish fastACCORD as a practical framework for ultrahigh-dimensional multi-omic partial correlation modeling and demonstrate how joint modeling of transcriptomic and epigenomic measurements can refine the interpretation of gene co-expression networks.

## Introduction

Partial correlation modeling is a principled approach to summarize residual linear associations among a large set of variables in multivariate data, enabling network-oriented interpretation. For variables with a well-defined positive definite covariance matrix, the partial correlation between two variables is the correlation between their residuals after linearly adjusting each for all remaining variables. Equivalently, it is obtained as the negative standardized off-diagonal entry of the inverse covariance matrix, or the precision matrix. Thus, sparse precision matrix estimation offers a natural framework for learning partial correlation networks from high-dimensional omics data, which can be further developed as a major tool to integrate complex multi-omic data.

When the variables are multivariate normal, zero partial correlation is equivalent to conditional independence, giving rise to the Gaussian graphical model interpretation (1). Currently, sparse estimation of such networks in high-dimensional gene expression data is commonly performed using methods such as the graphical lasso, which imposes an ℓ1 penalty on precision matrix entries (2,3). Related approaches, including sparse partial correlation estimation (SPACE) and constrained *l*_1_ -minimization for inverse matrix estimation (CLIME), also exploit sparsity in partial correlation or precision matrix structures, although they differ in objective functions and estimation principles (4,5).

In cancer transcriptomes, it is important to make the distinction between marginal co-expression and residual association, since the former may reflect indirect, shared, or confounded sources of variation whereas the latter may be an outcome of co-regulation or regulatory relationship. An RNA partial correlation network can be viewed as a conditional co-expression graph, in which an edge represents a transcript-to-transcript association not explained by the other transcripts included in the model (6–9). In this context, partial correlation estimators have previously been applied to gene expression data to prioritize prominent transcription factors (TFs), with potential extension to the identification of active gene regulatory network (GRN). Most success have come from simpler organisms or plants where TF-to-TF relationships are scarcely known due to the lack of experimental data supporting regulatory relationships in the form of direct DNA binding events and regulatory sequences (10–13).

In the context of modern GRN inference using the whole mammalian transcriptome data as input, partial correlation estimators are increasingly used to identify an undirected graph of all genes containing associations between TFs and their regulon, together with the relationships of other genes that are not fully explained by TF activity. Such a network may also capture associations arising from post-transcriptional regulation or RNA turnover. Hence the network estimated from full transcriptome-scale gene expression data is likely a broad superset of associations, within which direct TF-mediated regulatory relationships are embedded (2,5,14–22).

Conceptually, partial correlation networks are useful for the integration of multi-omic data when TF-mediated regulation and other molecular factors that modulate gene expression are measured in the same biological samples. TF-driven regulation and epigenetic regulation simultaneously contribute to transcriptome variation, and recent studies have highlighted the value of incorporating epigenomic measurements, such as chromatin accessibility (23,24) and DNA methylation (25), into GRN inference when such data are available alongside gene expression profiles. By the same token, in networks inferred using mRNA data only, covariation driven by unmodeled epigenomic features can be absorbed into apparent gene-gene co-expression and attributed to TF-associated co-regulation. By including measured epigenomic features in the same partial correlation model, joint multi-omic network estimation can unravel two complementary relationships: one representing epigenomic feature-gene dependencies that link regulatory marks to expression variation, and the other representing gene-to-gene associations that persist after measured epigenomic covariation is accounted for.

Despite this conceptual advantage, partial correlation network estimation has not been popularized for multi-omic analysis in the literature. This gap is primarily attributable to the unusual scale of the computational problem. A transcriptome-scale analysis already involves approximately 20,000 gene-level variables, and DNA methylation array data can add as many as 450,000 CpG measurements. In cohorts with multiple omic profiles, joint modeling is needed to estimate both epigenomic feature-to-gene dependencies and gene-to-gene associations adjusted for measured epigenomic covariation. The resulting sparse precision matrix estimation task calls for methods that exploit sparsity and avoid inversion of a general matrix defined over the full feature space. Consequently, graphical model-based multi-omic analyses in biomedical studies have typically been limited to reduced feature sets, such as fewer than 2,000 genes and a comparable number of CpG sites, rather than full transcriptome-methylome networks (8).

To overcome this barrier, we developed fastACCORD, a scalable extension of the ACCORD pseudolikelihood framework for ultrahigh-dimensional partial correlation modeling (26). fastACCORD combines an ℓ2-stabilized formulation for collinear multi-omic features, a semismooth Newton solver for efficient optimization, and row-separable computation that allows row blocks to be processed independently on a single CPU/GPU workstation without requiring an MPI-based distributed-memory HPC environment. Implemented in PyTorch, fastACCORD is designed to be maintainable and extensible across CPU and CUDA-enabled GPU environments, lowering technical barriers to genome-scale multi-omic network analysis.

We demonstrate fastACCORD in a pan-cancer integration of transcriptome and methylome profiles measured in the same primary tumor samples from 16 TCGA cancer types (27). For each cancer type, we estimated a joint partial correlation network containing more than 300,000 molecular features, enabling full-resolution analysis without pre-filtering the methylome to a small candidate set.

We used these networks to examine methylation-to-gene dependencies at promoter proximal and gene-body regions, compare methylation-adjusted gene-to-gene networks with mRNA-only networks, and test whether gene-to-gene associations retained after methylation adjustment are enriched for independent TF-associated regulatory evidence from ChIP-seq and curated regulon resources. We further compared TF-centered subnetworks across cancer types to characterize cancer-specific regulatory architectures and developed an interactive visualizer for exploring the resulting pan-cancer networks with ReMap ChIP-seq and JASPAR motif annotations (28,29).

Together, this work establishes fastACCORD as a scalable framework for ultrahigh-dimensional partial correlation modeling of multi-omic data and uses a pan-cancer transcriptome-methylome application to show the biological value of modeling multiple molecular layers jointly. The resulting networks provide a resource for studying methylation-expression coupling, methylation-adjusted co-expression, and TF-associated regulatory programs in cancer.

## Materials and Methods

### Data set selection and curation

All source data, including mRNA expression, microRNA expression and DNA methylation, were downloaded using TCGAbiolinks for 16 different cancers (30). These cancer types were selected based on two criteria: (i) DNA methylation was profiled using Illumina Human Methylation 450 platform and (ii) both RNA-sequencing data and DNA methylation array data are available for at least 100 primary tumor samples. Genomic coordinates of TSSs and CpG island locations were based on hg38. DNA methylation was quantified as beta values after Fisher transformation and mRNA and miRNA expression was quantified as log2(TPM+1). We only included genes and microRNAs if the TPM values were non-zero at least in 80% of the samples.

### Partial correlation network estimation with fastACCORD

Let *X* ∈ *R^n^*^×*p*^denote the centered and standardized data matrix, where *n* is the number of samples and *p* is the number of molecular features, and let **S** = n^−1^**X^T^X** be the sample covariance matrix. For a population covariance matrix Z with precision matrix *Θ* = (*θ_ij_*) = Σ^−1^, the partial correlation between variables *i* and *j* is 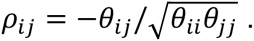 Thus, estimating a partial correlation network is equivalent to estimating the sparse off-diagonal structure of the precision matrix and standardizing its selected entries. This definition requires a well-defined positive definite covariance matrix; under multivariate normality, zero partial correlation is further equivalent to conditional independence, giving the usual Gaussian graphical model interpretation.

Conventional likelihood-based sparse precision matrix estimators, such as the graphical lasso, solve an optimization problem of the form

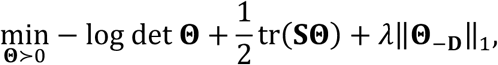

where *λ* > 0 is a regularization parameter, **Θ**_–**D**_ is a matrix without diagonal components of **Θ**, and **Θ** ≻ 0 denotes the positive definiteness of the symmetric matrix **Θ**. The gradient of this objective involves evaluating **Θ**^−1^, which is generally dense even when **Θ** is sparse. This matrix inversion operation becomes the main computational bottleneck when transcriptomic variables are modeled jointly with hundreds of thousands of epigenomic features (26).

fastACCORD instead builds on the ACCORD pseudolikelihood formulation (26). The method estimates a reparameterized precision structure

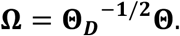

This reparameterization preserves the off-diagonal sparsity pattern of **Θ**. fastACCORD estimates **Ω** by finding the minimizer of the *P*_1_- and *P*_2_-regularized pseudolikelihood

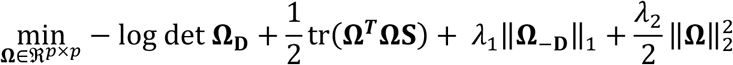

where **Ω_D_** and **Ω**_–**D**_ denote matrices only with and without diagonal parts of **Ω,** respectively. Here, the regularization parameter *λ*_1_ > 0 controls sparsity of the minimizer, and *λ*_2_ > 0 provides additional stabilization for highly collinear multi-omic features (31). The smooth part of this objective has gradient terms involving **ΩS**, **Ω**, and the diagonal matrix **Ω_D_**^−**1**^, but does not require the dense full inverse **Ω**^−**1**^ or **Θ**^−**1**^.

A key computational property of the ACCORD pseudolikelihood is row separability. The objective can be written as

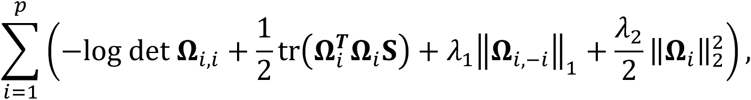

where **Ω***_i_* denotes the *i* -th row of **Ω**, **Ω***_i_*_,*i*_ and **Ω***_i_*_,–*i*_ denotes the diagonal and the off-diagonal entries in that row, respectively. Therefore, the optimization problem for the full *p* × *p* matrix variable can be decomposed into row-wise or row-block-wise sub-problems. fastACCORD solves these subproblems using a semismooth Newton method (32). The additional *l*_2_ term ensures positive definiteness of the Newton linear systems, improving numerical stability when active sets are large or when multi-omic features are strongly collinear. (Refer to **Supplementary Information** for details.) The solution 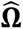 to the above optimization problem is the ACCORD estimator of **Ω.**

After estimating **Ω**, we reconstruct the precision matrix using the relationship

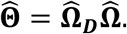

Because the row-wise optimization can yield small asymmetries in finite samples, we use the symmetrized estimate

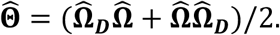

The estimated partial correlation between variables *i* and *j* is then computed as

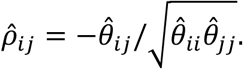

A network edge was recorded when the corresponding off-diagonal entry was selected as nonzero in the estimated sparse structure. For large multi-omic datasets, fastACCORD processes designated row blocks independently on CPU or CUDA-enabled GPU devices, allowing full or partial network estimation without storing or inverting general *p* × *p* matrices. Details of the semismooth Newton updates and implementation are provided in the **Supplementary Information**.

### Implementation and row-block execution

The row-wise decomposition of the fastACCORD objective allows the estimation task to be partitioned into row blocks. Each block can be assigned to a CPU or CUDA-enabled GPU process and solved independently, after which the selected rows are combined to form the full sparse network. This design avoids slow network-based communication (e.g. HP-ACCORD(26)) across compute nodes and allows users to adjust row-block size according to the feature dimension, expected graph sparsity, and available device memory.

### Selection of regularization parameters

Choosing an appropriate magnitude of *λ*_1_ is crucial for the performance of the computed estimator. At the multi-omic scale, we avoid the heavy computational burden of repetitive model retraining in cross-validation by adopting a computationally efficient BIC-based selection criterion. In particular, we used the extended pseudo-BIC (epBIC) (26,33), which is defined as

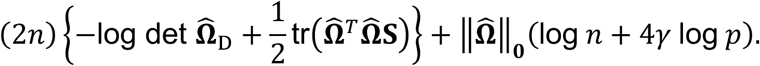

For all data, we selected the value of *λ*_1_ with the minimal epBIC from the candidate grid of 0.28, 0.31, … , 0.70 with fixed *λ*_2_ = 0.01 and *γ* = 0.3.

### Debiasing partial correlation estimates

Although the ACCORD estimator form effectively selects a graph underlying partial correlation network, the initially reported precision and partial correlations are biased estimates due to the shrinkage effect of regularization. For more accurate estimation, a debiasing procedure has been proposed (26). This procedure re-estimates 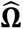 over the fixed support of the selected graph obtained during the selection phase, employing a smaller penalty to alleviate the bias in estimates. However, the debiasing procedure suffers from a drawback that the off-diagonal entries of 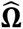 are inflated when the penalties are relaxed. As a consequence, for highly correlated variables, the magnitude of partial correlation estimates may exceed one, falling outside the feasible range. To navigate the trade-off between the underestimation inherence in biased estimation and infeasible overestimation in debiasing procedure, we propose the following refitted estimator 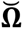 that minimizes the following objective function:

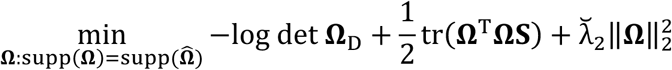

where the new hyperparameter 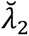 is chosen as an adequate value that the partial correlation derived from the final estimate 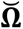 falls into the feasible range of |*ρ̂_i_*_j_| ≤ 1 while maintaining the support from the original estimator 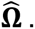 Empirical results show that imposing an equal or greater squared *l*_2_ norm in the penalty selection phase (*λ*_2_) exerts shrinkage effect stronger on artificially inflated estimates due to high correlation among the variables, whereas moderate estimates experience substantially less shrinkage. Therefore, we evaluated that the refitted estimator above successfully strikes a balance between alleviating the bias from graph selection phase and mitigating the inflation of the estimated 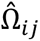 observed between highly correlated variables.

### Hardware requirements for TCGA analysis

fastACCORD offers options to compute ACCORD estimators on either CPU or CUDA-enabled GPU. For faster analysis of high-dimensional data with large *p*, using a CUDA device is highly recommended. Computation time decreases as the number of available devices increases. To avoid out-of-memory error, it is necessary to flexibly adjust the number of iterating rows handled in a single process based on the dimension *p* of the data, the expected sparsity of the estimator, and the available device memory. Empirically speaking, it is recommended to have at least 12GB of VRAM available in the CUDA device to run an iteration for 500 rows at once for data with *p* up to 300,000.

### TCGA network summary

For each cancer, all edges from the integrated network were assigned to one of three types (mRNA-mRNA, DNA methylation-mRNA, or mRNA-miRNA) based on node type annotations. For the mRNA-mRNA co-expression network comparisons, only edges with positive partial correlations were retained.

### M2G annotation and enrichment analysis

A M2G edge was classified as cis if the DNA methylation probe and its paired gene were on the same chromosome, within 1 Mb, and the probe was annotated as regulating that specific gene in the Illumina 450K array metadata. Among cis edges, probe genomic location was annotated as promoter proximal (TSS1500, TSS200, 5’UTR, 1st Exon) or gene body (Body, 3’UTR). Cis edges with negative partial correlation at promoter proximal probes were interpreted as methylation-associated silencing, and those with positive partial correlation at gene body probes as methylation-associated activation.

For each cancer, KEGG pathway enrichment was performed separately on three gene sets defined by cis methylation direction and probe location: (1) promoter proximal probes (TSS1500, TSS200, 5’UTR, 1st Exon) with negative partial correlation (promoter-silenced); (2) gene body probes (Body, 3’UTR) with positive partial correlation (gene-body-activated); and (3) gene body probes with negative partial correlation (gene-body-suppressed). For the pan-cancer bubble plot, pathways were retained if they were significantly enriched (p < 0.05) in at least 3 cancers in either the promoter-silenced or the gene-body-suppressed sets (union). Pathways from the Human Diseases KEGG category and non-informative subcategories were excluded, and remaining pathways were grouped into four broad classes: Metabolism, Signalling, Immune, and Others.

Genes with cis promoter methylation and negative partial correlation in at least three cancer types were defined as commonly methylation-silenced. GO Biological Process and KEGG pathway enrichment were performed on this gene set using clusterProfiler, with all genes observed in the methylation-mRNA network as the background universe and BH-adjusted p-values.

### ReMap ChIP-seq data curation

ChIP-seq data were obtained from the ReMap 2022 database (28), specifically the non-redundant (NR) peak catalog (remap2022_nr_macs2_hg38_v1_0.bed, hg38), which consolidates ChIP-seq experiments targeting the same TF across cell types into consensus peaks. Each NR peak encodes the contributing TF and all cell types in which the peak was detected. Candidate TF binding sites were defined as NR peaks whose midpoint fell within ±500 bp of the TSS of each protein-coding gene in the ENSEMBL GRCh38 v115 annotation, where the TSS was taken as the 5′ end of the gene model on the annotated strand. A network edge was labelled as a TF-target relationship if one gene in the pair encoded a TF with at least one ChIP-seq peak within ±500 bp of the other gene’s TSS.

To validate TF binding events in a cancer-specific context, for each of the cancer types, cell lines and tissues with ChIP-seq data in ReMap NR were manually selected based on biological relevance, and TF-target annotations were restricted to peaks detected in those cancer-relevant contexts. TF-target subnetworks were visualized using NetworkX and Matplotlib in Python.

### DepMap resource

CRISPR gene essentiality data were obtained from the DepMap portal (release 26Q1) (34), using Chronos gene effect scores (35) from the CRISPRGeneEffect.csv file and cell line metadata from Model.csv. For each of the 16 cancer types, primary tumor cell lines were selected and matched to the corresponding TCGA cohort via OncotreeCode mapping. Mean Chronos scores were computed per gene per cancer type, and genes with a mean Chronos score ≤ −0.5 (or ≤ −1.0 for stringent essentiality) were classified as essential.

### TF subnetwork

For a comprehensive TF annotation, we used the list of >1600 human TFs reported by Lamber et al (36). TF family classification was applied to TF subnetwork analysis. For each TF, a two-layer subnetwork size was computed per cancer type using TF-gated propagation: Layer 1 comprises all direct neighbors of the TF, and Layer 2 extends through Layer-1 nodes that are themselves TFs. Subnetwork specificity was quantified using the tau (*τ*) statistic applied to the subnetwork size across all 16 cancer types. For each TF, let *x_i_* denote its subnetwork size (excluding the root TF itself) in cancer *i* and *N* is the number of cancers. The statistic is defined as 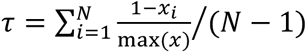 (37), yielding values near 1 for cancer-specific TFs and near 0 for pan-cancer activity. An analogous *τ* score was computed for expression specificity using mean tumor TPM across cancers. To assess the concordance between where a TF is most network-active and most highly expressed, both subnetwork size and expression values were row-standardized to z-scores across cancer types per TF across all (TF, cancer) pairs.

### G2G network visualizer

We developed an interactive web-based visualizer for exploring methylation-adjusted G2G networks across TCGA cancer cohorts. Users can query one or multiple genes (up to 10) and visualize their subnetworks in one or multiple cancer types (up to 8). A shortest path option supports visualization of connections among up to 50 selected genes. Users can also search for TFs, select TFs by family, and explore their TF-gated subnetworks. Multiple edge-coloring options are provided to annotated G2G relationships using ReMap ChIP-seq evidence (as discussed in the results), JASPAR motif evidence, their union, or cancer cohort membership.

For JASPAR motif-matching annotation, TF binding motifs were obtained from JASPAR CORE vertebrate collection using R packages JASPAR2024 (the latest version available in Bioconductor) and TFBSTools (29). JASPAR motif reference was built for all human protein-coding genes with hg38 as reference genome. The TSS ±500 bp promoter region was scanned for matches to JASPAR position-weight matrices using motifmatch with a significance threshold of p-value < 10^-4^, generating a binary gene-by-motif matrix. To annotate edges in the network, JASPAR TFs with mean expression greater than 1 TPM were retained. JASPAR-derived TF-target and co-regulating pairs were annotated the same way as in ReMap.

## Results

### fastACCORD makes ultrahigh-dimensional multi-omic partial correlation modeling tractable

Estimating joint transcriptome-to-methylome partial correlation networks requires sparse precision matrix estimation over hundreds of thousands of molecular features. Conventional Gaussian likelihood-based methods, including graphical lasso (2) and BigQUIC (38,39), depend on operations involving the dense inverse of the precision matrix iterate, creating a major computational bottleneck when hundreds of thousands of epigenomic features are modeled jointly with transcriptomic variables. fastACCORD avoids this bottleneck by building on the ACCORD pseudolikelihood formulation (26,40), whose gradient requires no matrix inversion and whose objective is separable across rows. We further incorporate an *P*_2_-stabilized objective to improve numerical stability for highly collinear multi-omic features and use a semismooth Newton solver to accelerate row-wise optimization (**Figure 1A**).

**Figure 1.**
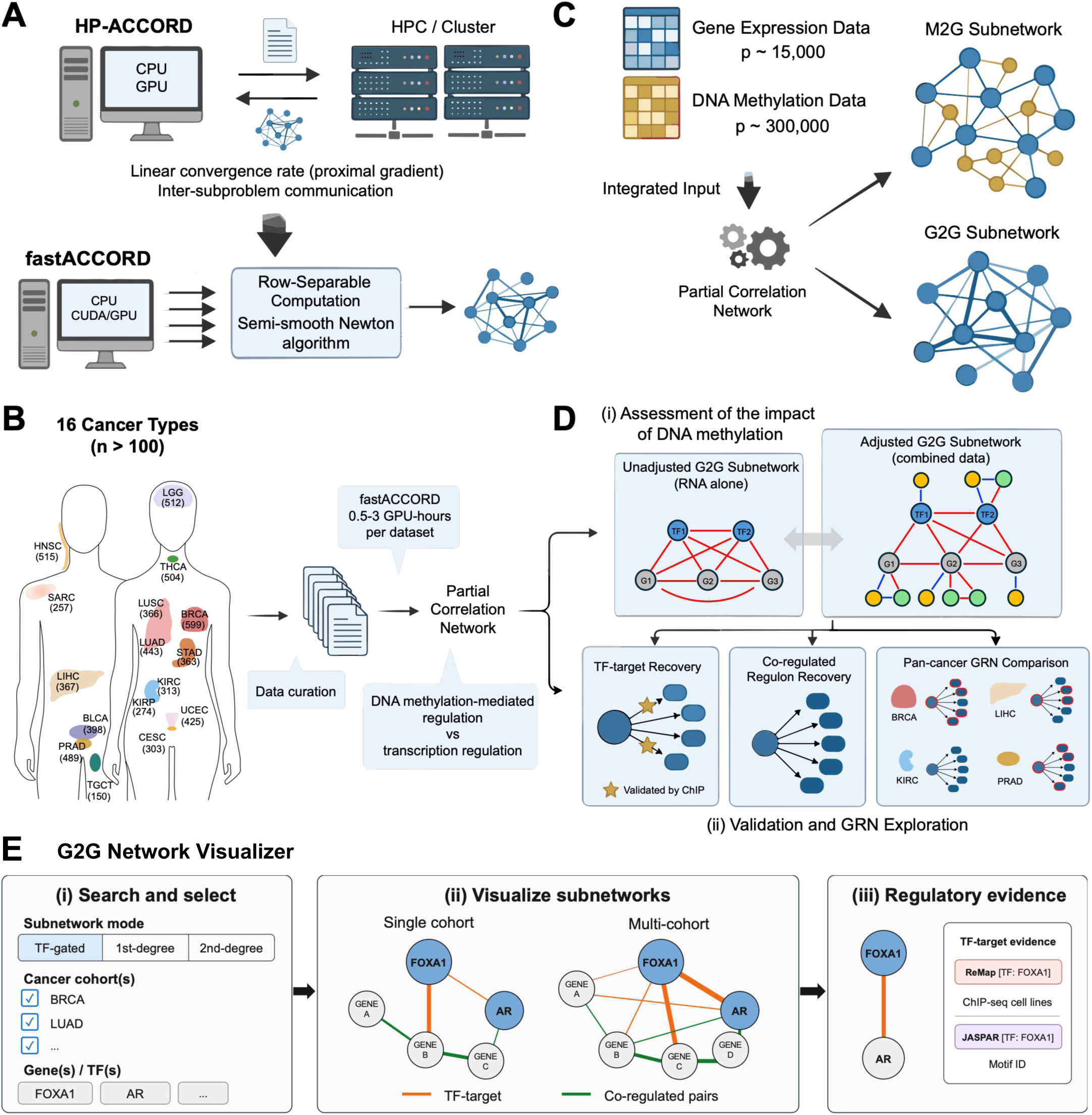
(**A**) Schematic overview of the transition of fastACCORD from HP-ACCORD, including development of a faster algorithm and implementation of row-separable computation for greater applicability on standard hardware. (**B**) Partial correlation network analysis of gene expression and DNA methylation data using fastACCORD in sixteen cancer types of TCGA. (**C**) Concept of methylation-expression subnetwork (M2G) and gene-to-gene (G2G) subnetwork derived from partial correlation network. Methylation-methylation subnetwork (M2M) was not interpreted in this work. (**D**) Comparison of G2G subnetworks with and without adjustment for DNA methylation data, called adjusted analysis (integrated data) and unadjusted analysis (mRNA data alone) respectively. (**E**) Interactive network visualizer. Users can search multiple genes and cancer types, visualize individual or pan-cancer interaction networks, and inspect edge annotations with potential regulatory relationships. **Alt Text**: Overview of fastACCORD and its pan-cancer application. Optimized computational framework enables large-scale partial correlation network estimation on standard CPU and CUDA-enabled GPU hardware. Matched DNA methylation and RNA-seq gene expression data from 16 TCGA cancer types are integrated using fastACCORD. The integrated networks are evaluated for TF-associated regulatory evidence and explored through an interactive network visualizer.

This row-separable design allows designated row blocks to be processed independently on CPU or CUDA-enabled GPU devices, avoiding the need for a HPC environment (26). The PyTorch implementation is designed to be maintainable and extensible across common computing environments, lowering software and hardware barriers for genome-scale network analysis. Because fastACCORD estimates row blocks without storing or inverting unstructured, high-dimensional matrices, its practical memory requirement is governed primarily by row-block size and active-set cardinality of the iterator.

Benchmark experiments confirmed that these design choices improved scalability at multi-omic feature dimensions. Across synthetic and TCGA-derived tasks, fastACCORD scaled to up to a million variables, where the row-block size and sparsity regularization parameter was configured to ensure the rapid computation within the memory constraint of the available CUDA device. For datasets with a reasonable number of features where direct comparison was feasible, fastACCORD exhibited faster and more memory-efficient computation than conventional sparse precision estimation methods, while maintaining support recovery accuracy in simulated sparse partial correlation networks. For example, for Gaussian samples with a sparse precision matrix with n=100 and p=10,000, fastACCORD required approximately 50 and 65 seconds to obtain a single precision matrix estimate using a single GPU and CPU device, respectively, whereas the official CPU implementations of QUIC and CLIME required approximately 70 and 500 seconds, respectively. Since fastACCORD is row-separable, further reductions in computation time can be achieved by distributing the row-wise optimization task across multiple GPU devices without inter-device communication.

These results establish fastACCORD as a practical framework for full-resolution multi-omic partial correlation network estimation and motivate its application to pan-cancer transcriptome-methylome network analysis.

### Pan-cancer construction of joint transcriptome-methylome networks

We next applied fastACCORD to TCGA data comprising 16 cancer types (27), using mRNA expression and DNA methylation profiles measured in the same primary tumor samples (**Figure 1B** and **Table S1**). Cohorts were selected using three criteria: (i) samples were primary tumors, (ii) each cohort contained at least 100 primary tumor samples, and (iii) both RNA-seq expression data and Illumina HumanMethylation450 DNA methylation data were available for the same samples (see **Materials and Methods**). After feature curation, each cancer-specific data matrix contained protein-coding gene expression, miRNA expression and CpG methylation features, yielding more than 300,000 molecular variables per cancer type (**Table S1**). For each cohort, we estimated a joint transcriptome-methylome partial correlation network using fastACCORD. In parallel, we estimated an mRNA-only partial correlation network from the same tumor samples and protein coding genes. Although miRNA expression data was included in the full integrated network, the total number of miRNA species was limited and we largely focused on methylation-to-gene and gene-to-gene subnetworks and omitted the discussion of miRNAs in this work.

We decomposed the resulting networks into edge type-specific subnetworks. We define the methylation-to-gene (M2G) subnetwork as the DNA methylation-mRNA component of the joint network, representing dependencies between CpG methylation features and mRNA expression after accounting for all other measured molecular variables. We also define the methylation-adjusted gene-to-gene (G2G) network as the mRNA-mRNA component of the joint network, in which transcript-to-transcript residual associations are estimated after accounting for measured DNA methylation covariation. By contrast, the unadjusted G2G network was defined as the mRNA-mRNA partial correlation network estimated from the corresponding mRNA expression data alone.

Both joint and RNA-alone networks were estimated using fastACCORD on a single multi-GPU workstation using up to four NVIDIA TITAN V GPUs with 12 GB VRAM each. Across cancer types, obtaining a selected fastACCORD estimate required approximately 0.5-3 GPU hours, depending on the number of molecular features and the density of the estimated graph. These computational results demonstrate the feasibility of full-resolution pan-cancer transcriptome-methylome network construction reducing reliance on HPC environments, which are typically less accessible and more operationally complex than commodity GPU hardware. Additional description of workflows and computational performance are available in **Supplementary Information**.

Using the joint and mRNA-only networks, we interpreted the pan-cancer results around the following two main questions. First, to what degree is DNA methylation at upstream regulatory and gene body coupled to gene expression? Second, do methylation-adjusted G2G networks capture shared TF-associated co-regulation with greater specificity than mRNA-only networks? Both questions require ascertainment of residual associations among CpG methylation features and gene expression in an ultrahigh-dimensional model, which is enabled by fastACCORD.

Specifically, we evaluated each cancer-specific partial correlation network from several angles. First, we examined M2G subnetworks and compared the adjusted G2G subnetwork estimated from the integrated data and the unadjusted G2G subnetwork estimated from the gene expression data alone (**Figure 1C**, **1D**). In doing so, we assessed how measured DNA methylation covariation modulates co-expression networks of surrounding genes across the cancers (**Figure 1C**). Second, we assessed the recovery rates of known TF-target relationships embedded in the partial correlation networks and the recovery of co-regulated targets of shared TFs (**Figure 1D)**. Third, we used the resulting methylation-adjusted G2G subnetwork to compare TF-associated network structures across the 16 cancer types and investigated whether multiple cancers share TF regulation patterns.

Based on these results, we also developed a network visualizer which allows users to navigate the networks for prioritized TF genes with cancer-specific co-regulatory architectures (**Figure 1E**). The visualizer supports searches for multiple genes in one or more cancer cohorts and annotates network edges with regulatory evidence from the ReMap ChIP-seq resource and JASPAR motif matches (28,29). To our knowledge, this work provides the first pan-cancer genome-scale transcriptome-methylome network resources to compare TF-associated gene-to-gene architectures in a way that accounts for the modulatory role of DNA methylation covariation in shaping gene co-expression. The resource offers a practical foundation for prioritizing cancer-specific TFs and exploring their co-regulatory landscapes.

### Joint networks reveal a pan-cancer landscape of methylation-expression dependencies

The joint transcriptome-methylome networks first allowed us to examine M2G dependencies across cancers. As shown in **Figure S1** and **Figure 2A**, G2G edges were captured predominantly as positive partial correlations, whereas M2G edges featured both positive and negative directions. Because local methylation-expression relationships have the most direct biological interpretation, we focused on cis-M2G association, defined as the edge linking a DNA methylation site to the nearby gene on the same chromosome, where the CpG feature was previously annotated to the gene and located in either an upstream regulatory region or the gene body (see **Methods**).

**Figure 2.**
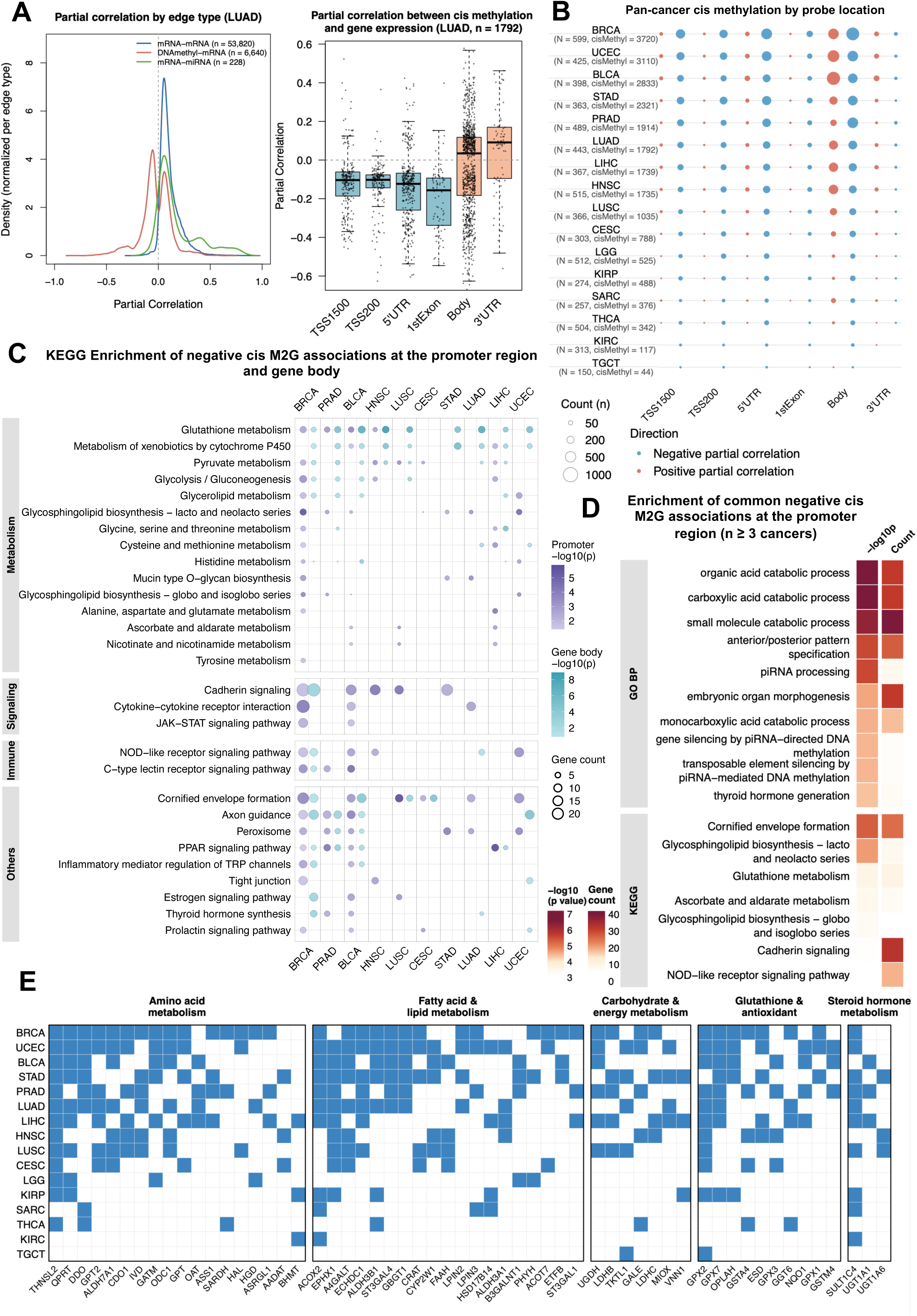
Methylation-to-gene (M2G) edges in the multi omics network across cancers. (A) Representative kernel density plot of partial correlation by edge type (mRNA-mRNA, DNA methylation-mRNA, mRNA-miRNA; left) and partial correlation between cis methylation and gene expression stratified by probe location in LUAD (right). (B) Cis-M2G edge counts by probe location and direction. N: cohort size; cisMethyl: total cis-M2G edges per cancer. (C) KEGG pathway enrichment of genes with negative cis-M2G associations at promoter and gene body regions across cancers. (D) KEGG and GO BP enrichment of genes recurrently silenced by promoter cis methylation in ≥3 cancers. (E) Metabolic gene membership among promoter cis methylation-silenced genes in ≥3 cancers. **Alt Text**: Pan-cancer methylation-to-expression relationships inferred from joint networks. Promoter proximal methylation is predominantly negatively associated with gene expression, whereas gene body methylation shows both positive and negative associations. Recurrently observed promoter region methylation-associated expression repression events (negative partial correlation) are enriched for metabolic genes across multiple cancer types.

The extent of cis-M2G connectivity varied markedly across cancers. BRCA, UCEC, BLCA and STAD showed the most extensive cis-M2G connections, whereas THCA, KIRC and TGCT showed relatively few, suggesting that tissue- or cancer-specific epigenetic dysregulation contributes to the variation in M2G connectivity beyond the differences in cohort size (**Figure 2B**). The sign of cis-M2G associations depended strongly on genomic location. Promoter methylation sites proximal to the transcription start site (TSS), including TSS1500, TSS200, 5′UTR and 1stExon annotations, were predominantly negatively associated with expression, consistent with promoter methylation-associated repression. By contrast, gene-body methylation past the first exon showed both positive and negative partial correlations, consistent with a mixture of positive associations reflecting active transcription and negative associations reflecting hypermethylation-induced suppression.

We next examined which biological programs were represented by these local methylation-expression dependencies. Genes connected to promoter proximal CpG features by negative cis-M2G edges were enriched for processes including cornified envelope formation (notably in LUSC), cadherin signalling, glutathione metabolism and central metabolic pathways (**Figure 2C** and **Figure S2**). Genes connected to gene body CpG features by negative cis-M2G edges showed a similar enrichment profile, likely reflecting hypermethylation encroachment beyond promoter CpG islands into flanking gene body regions. In these cases, promoter proximal and gene body CpG features may capture the same methylation-associated silencing event from different genomic positions (41). In contrast, genes positively associated with gene body methylation were consistently enriched for signalling pathways such as calcium and cAMP signalling across cancers (**Figure S3**), consistent with the canonical role of gene body methylation as a marker of transcriptional activity in these cases (42,43).

To identify methylation-repressed gene expression programs shared across cancers, we focused on genes with negative cis-M2G associations at promoter proximal regions in three or more cancer types. KEGG and GO enrichment of this recurrent gene set identified small molecule metabolism as the most significantly enriched category (**Figure 2D** and **Table S2**), including amino acid catabolism (THNSL2, DDO, GPT/GPT2, CDO1 and QPRT), fatty acid oxidation and lipid metabolism (ACOX2, ACOT7, CRAT, PHYH, ECHDC1 and LPIN2/3), and carbohydrate metabolism (LDHB, LDHC and TKTL1; **Figure 2E**). Among these genes, THNSL2 showed the broadest recurrence, with negative promoter proximal M2G associations across 13 cancer types, raising the possibility that methylation-associated repression of THNSL2 may confer metabolic advantages supporting tumor growth. Additionally, LDHB showed negative promoter proximal M2G associations in five cancers, linking Warburg-associated metabolic reprogramming to methylation-associated repression of LDHB in a subset of tumor types beyond the canonical LDHA-driven pathway (44).

### Methylation adjustment reshapes co-expression network

Having characterized methylation-expression dependencies, we investigated how measured DNA methylation covariation alters the gene-to-gene component of the transcriptome network. We compared methylation-adjusted G2G networks, defined as the mRNA-mRNA components of the joint transcriptome-methylome networks, to unadjusted G2G networks estimated from mRNA expression data alone using the same tumor samples and protein-coding genes. For gene co-expression network comparison, we focused on positive gene-to-gene partial correlations (see **Methods**).

Across all cancer types, methylation-adjusted G2G networks were sparser than their unadjusted RNA-only counterparts, although the magnitude of reduction varied by cancer type (**Figure 3A** and **Table S1**). Cohorts with smaller sample sizes, including SARC, CESC, KIRC and KIRP, tended to lose more G2G edges after adjustment, consistent with reduced power to support stable edges in high-dimensional partial correlation models. However, edge reduction was not solely explained by cohort size. Several larger cohorts also showed substantial reductions in G2G connectivity after adjustment for dominant hypermethylation-associated co-expression structure in these tumors. Examples include LGG (45), where IDH-driven hypermethylation is a prominent molecular feature, and THCA, where BRAF V600E-associated hypermethylation has been reported (46). By contrast, cancers often associated with molecular heterogeneity, including BRCA, PRAD, UCEC and STAD, showed relatively modest edge reduction, possibly because divergent methylation patterns across subtypes limit the contribution of shared methylation covariation to cohort-wide co-expression.

**Figure 3.**
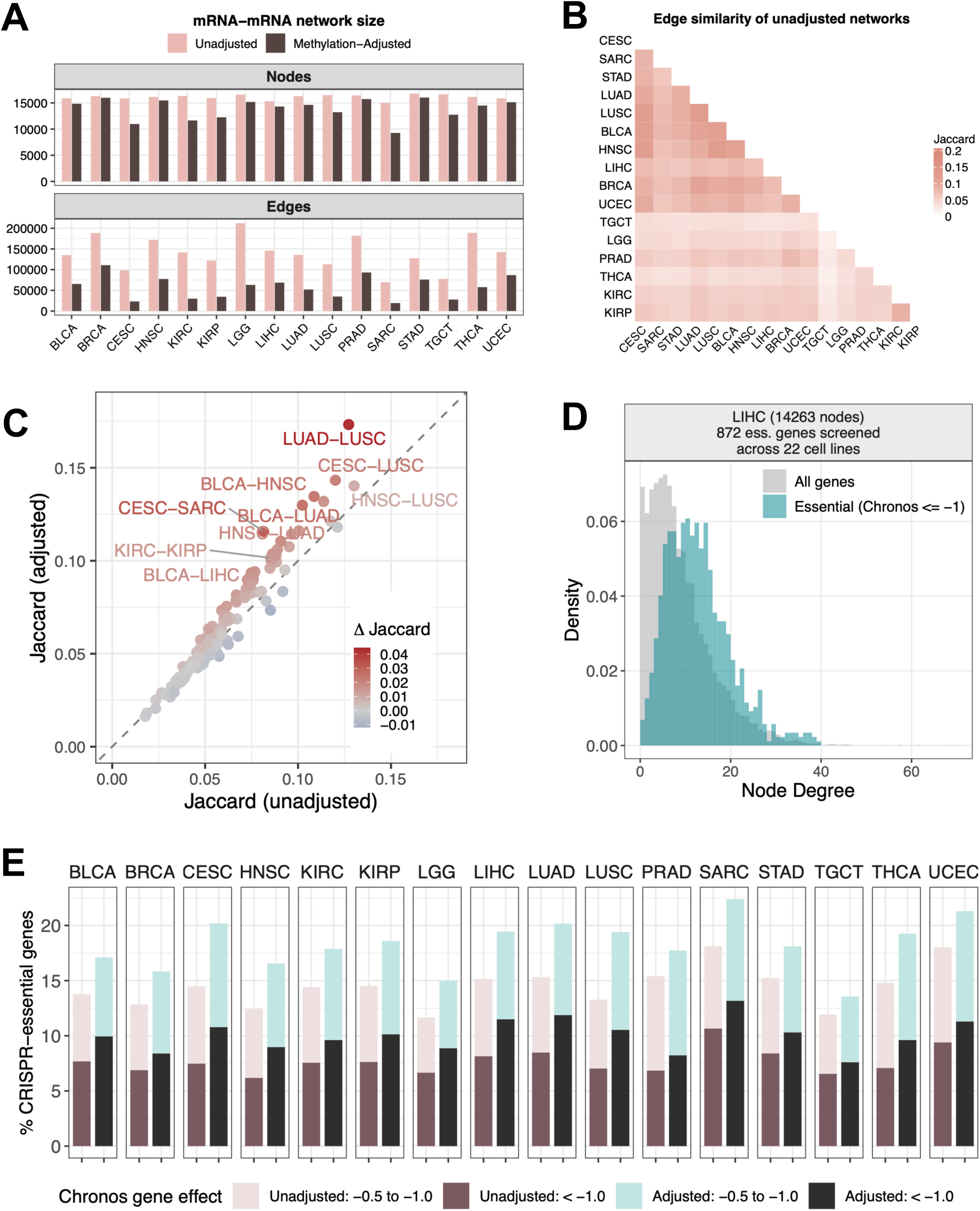
Gene co-expression network before and after methylation adjustment. (A) Size of the gene co-expression networks across cancers. (B) Edge similarity among methylation-unadjusted G2G networks, measured using the Jaccard similarity index. (C) Changes in Jaccard similarity after methylation adjustment. (D) Distribution of node degrees for all genes (gray) and CRISPR-essential genes (Chronos score < −1, blue) within the networks. LIHC is shown as an example. (E) Proportion of CRISPR-essential genes (Chronos < −1, dark; −0.5 to −1, light) among the top 20% highest-degree hub genes in adjusted and unadjusted networks across 16 cancer types. **Alt Text**: Comparison of gene-to-gene networks before and after adjustment for DNA methylation across the cancers. Methylation-adjusted networks contain fewer edges, show increased cross-cancer network similarity, and have high-degree hubs more enriched for CRISPR-essential genes than corresponding networks derived without DNA methylation adjustment.

We next examined how methylation adjustment affected cross-cancer similarity of G2G networks. In unadjusted mRNA-only networks, pairwise Jaccard similarity of edge sets revealed clustering by tissue of origin (KIRC-KIRP; LUAD-LUSC) and histological subtype (squamous cell carcinomas: BLCA, LUSC, CESC and HNSC) (**Figure 3B**). After methylation adjustment, between-cancer similarity increased across all cancer pairs (**Figure 3C**), suggesting that measured methylation covariation can obscure shared gene-to-gene architecture in the mRNA-only networks. The largest gain in similarity was observed between LUAD and LUSC, consistent with their distinct methylation landscapes that may mask shared regulatory structure in unadjusted networks (47,48). Hub genes contributing to the increased LUAD-LUSC similarity included immune cell markers (CD79B, JAML, CD2, PLEK) and housekeeping regulators of mitochondrial metabolism and transcription (**Figure S4**), reflecting a shared immune microenvironment (49) and core cellular programs at differentially methylated CpG-rich promoters (50). Overall, these analyses suggest that methylation adjustment can reveal shared gene-to-gene co-expression architecture that may be masked by cancer-specific methylation patterns.

Finally, we asked whether hub genes in methylation-adjusted G2G networks were enriched for genes functionally important for tumor-cell survival. We mapped CRISPR gene dependency scores (Chronos) from cancer type-matched DepMap cell lines onto network nodes (34,35). Within adjusted networks, CRISPR-essential genes (Chronos ≤ −1) were enriched among high degree nodes relative to all genes (**Figure 3D** and **Figure S5**). Moreover, the top 20% highest degree genes in adjusted networks contained a higher proportion of CRISPR-essential genes than the corresponding hubs in unadjusted networks across cancer types (**Figure 3E**).

Together, these results show that high degree nodes in methylation-adjusted G2G networks are more enriched for independent CRISPR-based essentiality signals than corresponding hubs in mRNA-only networks.

### Methylation-adjusted G2G networks are enriched for regulatory evidence associated with TF(s)

To investigate whether the methylation-adjusted networks aligned with external evidence of TF-associated regulation, we first mapped each gene-to-gene edge to the ReMap database, a TF-target database using ChIP-seq binding evidence (28). We used TF-binding annotations as orthogonal regulatory evidence to support the biological interpretation of the undirected G2G networks, rather than inferring regulatory direction. An edge was annotated as a TF-target relationship if one endpoint encoded a TF with at least one ReMap ChIP-seq peak near the TSS of the other endpoint gene (see **Methods**).

Across the 16 cancer types, methylation-adjusted G2G networks consistently showed a higher proportion of TF-target annotated edges than unadjusted networks (mean 6.54% vs. 5.39%; **Figure 4A**), with the gain ranging from 0.49% in STAD to 2.20% in KIRP. This pattern was also observed using cancer type-specific ReMap ChIP-seq backgrounds constructed from matched cell lines and tissues (**Table S4**, **Figure S6**). Notably, the higher annotation rate was observed even though methylation adjustment removed 39-79% of G2G edges across cancer types, suggesting that G2G edges retained after accounting for measured DNA methylation covariation are more concentrated among gene pairs supported by TF-binding evidence.

**Figure 4.**
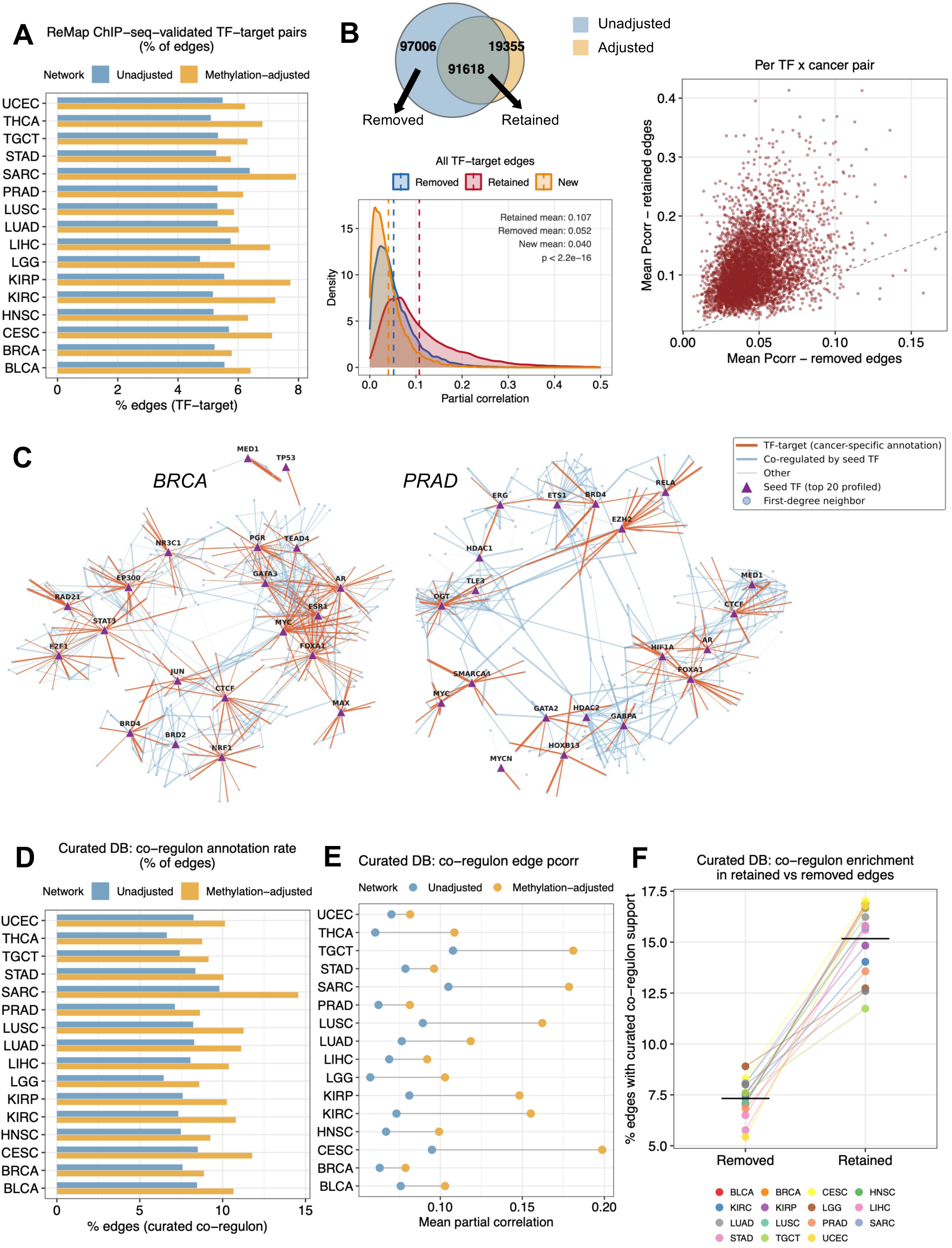
TF-target and co-regulatory annotation in methylation-adjusted and unadjusted gene co-expression networks. (A) Percentage of gene co-expression edges supported by ChIP-seq-validated TF-target interactions from ReMap, comparing unadjusted and methylation-adjusted networks across 16 cancer types. (B) Quality of TF-target edges retained or removed by methylation adjustment. The Venn diagram (BRCA shown as example) illustrates edge overlap between networks: edges unique to the unadjusted network (removed, blue), shared edges (retained, overlap), and edges unique to the adjusted network (yellow). The density plot shows partial correlations of retained versus removed TF-target edges; the scatter plot shows mean partial correlations of retained and removed edges per TF-cancer pair. (C) Co-regulatory subnetworks of the most ChIP-seq profiled TFs in cancer type-matched cell lines from ReMap, showing direct connections among these TFs within the adjusted network (BRCA and PRAD shown as examples). (D) Percentage of gene co-expression edges connecting gene pairs sharing at least one common upstream TF in an in-house curated TF-target database, comparing unadjusted and adjusted networks. (E) Mean partial correlations of curated co-regulon edges in unadjusted and methylation-adjusted networks per cancer type. (F) Proportion of curated co-regulon edges among edges retained in versus removed from the adjusted network across cancer types. **Alt Text**: Regulatory evidence supporting gene-to-gene networks before and after methylation adjustment. Methylation-adjusted networks have higher proportions of ReMap ChIP-seq-supported TF-target edges and curated co-regulon pairs. TF-associated edges retained after methylation adjustment have stronger partial correlations and greater co-regulon support than the removed edges, with representative TF-centered networks shown for breast and prostate cancer.

We then compared ReMap-annotated TF-target edges retained after methylation adjustment with those present only in the unadjusted mRNA-only networks. Retained TF-target edges carried substantially larger absolute partial correlations than those removed (**Figure 4B**), indicating that methylation adjustment preferentially retained higher-magnitude TF-associated gene-gene associations. Among the retained edges, TFs with the most extensive ChIP-seq profiling records in cancer type-matched cell lines (**Table S5**) were highly interconnected, forming co-expression hubs within the adjusted networks (**Figure 4C**). In BRCA, for example, ESR1, AR, MYC, FOXA1 and GATA3 were extensively interconnected, consistent with known hormone- and lineage-associated regulatory programs in breast cancer.

Beyond direct TF-target annotations, we examined whether G2G edges captured putatively co-regulated gene pairs sharing upstream TFs. More than 75% of G2G connections were supported by ReMap ChIP-seq evidence of at least three shared upstream TFs (**Figure S7**). This proportion was consistently higher in methylation-adjusted networks than in unadjusted mRNA-only networks, supporting the interpretation that methylation adjustment further enriches the connections supported by shared TF binding.

We further validated co-regulon support for G2G edges using a curated TF-target database (51) of over 10,000 human TF-regulon pairs compiled from TRRUST, ENCODE, TRED and ITFP (52–55). The adjusted networks contained a higher percentage of edges connecting gene pairs annotated as co-regulon pairs across all cancer types (**Figure 4D**). These co-regulon edges also showed higher average partial correlations in methylation-adjusted networks than in unadjusted networks (**Figure 4E**). Compared with the edges removed by methylation adjustment, the retained edges showed approximately two-fold higher co-regulon annotation rates (**Figure 4F**). Overall, these results indicate that accounting for DNA methylation covariation sharpens the TF-associated component of G2G co-expression networks, enriching the retained edge sets for independently supported TF-target and co-regulon relationships.

### TF-centered subnetworks reveal cancer-specific co-expression architecture

To characterize cancer-specific co-expression patterns around individual TFs, we extracted a two-layer subnetwork centered on each TF from the methylation-adjusted G2G network. Layer 1 comprised all direct co-expression neighbors of the TF, and Layer 2 comprised second-degree nodes reached through Layer 1 nodes that were themselves TFs (see **Methods**). As illustrated by FOXA1 (**Figure 5A**), the resulting TF-centered subnetwork size varied substantially across cancer types, indicating that the breadth of TF-associated co-expression connectivity is highly cancer specific.

**Figure 5.**
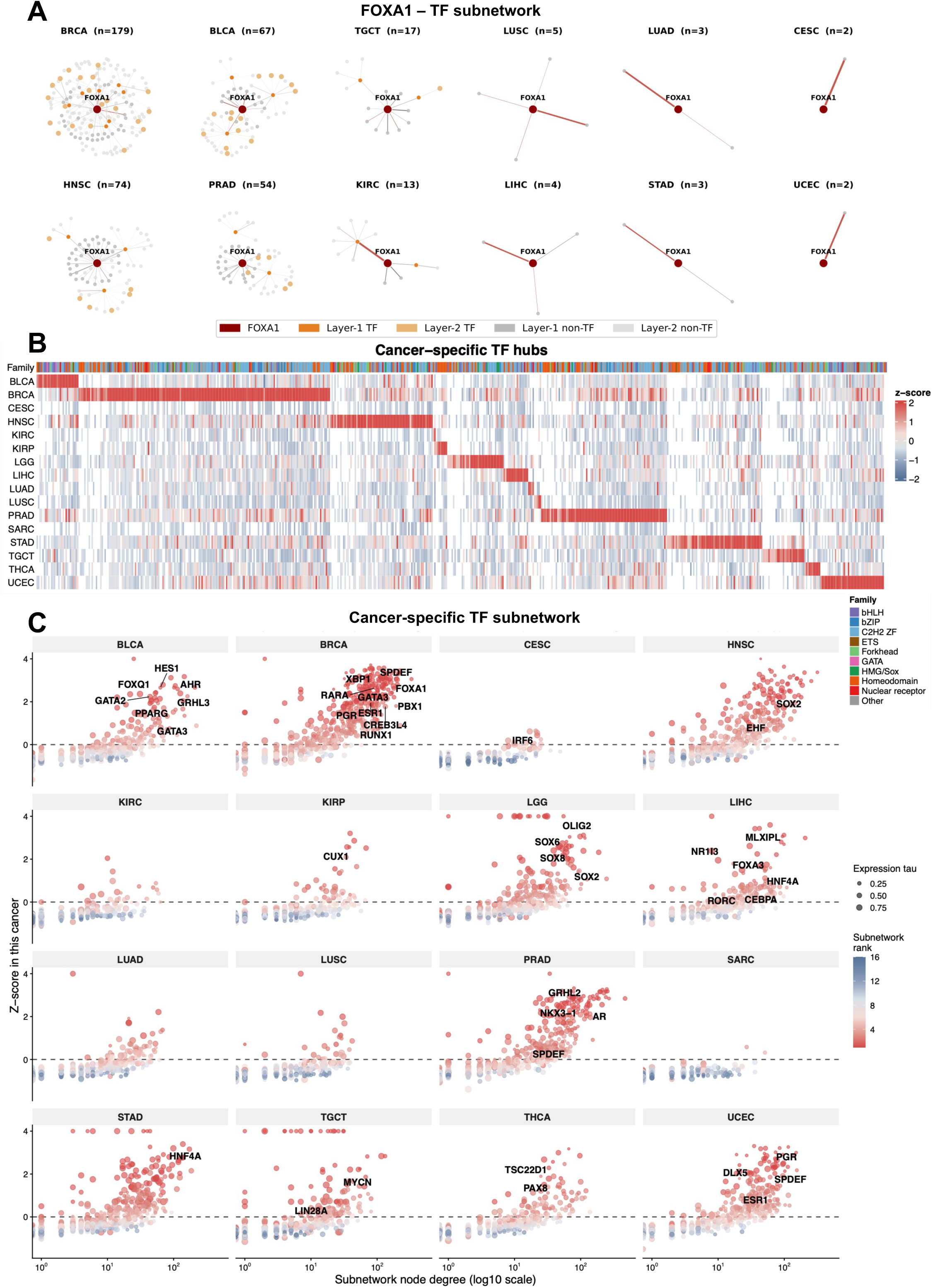
Cancer-specific TF co-expression subnetworks across 16 TCGA cancer types. (A) Representative two-layer TF subnetwork of FOXA1 in BRCA, illustrating the first-degree co-expression neighbors and second-degree nodes connected exclusively through first-degree TF nodes. (B) Heatmap showing the row z-score of subnetwork sizes for TFs with cancer-specific co-expression patterns (*τ* specificity score > 0.8, present in ≥4 cancer types) across 16 cancer types. TFs are ordered by the cancer in which they achieve their peak z-score. (C) Scatter plots showing subnetwork size (log10 scale) against z-score for each TF in each cancer type. Dot color reflects the cancer-type rank of each TF’s subnetwork size and dot size reflects expression *τ* specificity. TFs with established roles or recognized as master regulators in the corresponding cancer type (Table S7) are labelled where z-score > 0.5. Single-cancer exclusive TFs (*τ* = 1) are included and plotted at z-score = 4. **Alt Text**: Cancer-specific TF-centered view of co-expression architectures in methylation-adjusted networks. FOXA1 illustrates marked differences in subnetwork size across the cancers. Pan-cancer comparison of approximately 1,600 TFs identifies more than 650 with highly cancer-specific subnetworks, including known lineage-associated regulators such as ESR1 and FOXA1 in breast cancer, AR in prostate cancer, and HMG/Sox-family TFs in glioma.

To quantify the cancer specificity of each TF-centered subnetwork, we computed tau (*τ*) specificity scores for the two-layer subnetwork size across the 16 cancer types. Among ∼1,600 annotated human TFs (36), over 650 TFs spanning diverse TF families exhibited *τ* >0.8 (**Figure 5B**, **Table S6**), indicating that a substantial proportion of human TFs show highly cancer-specific regulon co-expression profiles. Several cancer types including SARC, CESC, KIRC and KIRP showed relatively few TFs with positive subnetwork-size z-scores (**Figure 5B**), suggesting that cancer-specific TF-centered connectivity was more difficult to isolate in these cohorts. This pattern may reflect reduced G2G network density after methylation adjustment (**Figure 3A**) and less coherent gene co-expression due to molecular heterogeneity of these cancer types, e.g. in SARC which encompasses multiple histologically and genomically distinct soft-tissue sarcoma subtypes (56).

We next compared these cancer-specific TF-centered subnetworks with a curated set of TF-cancer associations, comprising established cancer TFs and master regulators ranked by CaCTS scores (57) **(Table S7**). Many TFs with high z-scores of subnetwork size in a given cancer type corresponded to known master regulators in cancers (**Figure 5C**, labelled in the plots). At the TF family level, nuclear receptor TFs were prominent in hormone-driven cancers, including ESR1 in BRCA and AR in PRAD, reflecting estrogen receptor-dependent regulation in breast cancer and androgen receptor-driven prostate lineage specification, respectively (58,59). Forkhead TFs were prominent in BRCA, including FOXA1 (60), and in HNSC (61,62). HMG/Sox family TFs characterized the LGG block, consistent with the neural lineage context of glioma (63), whereas homeodomain TFs were enriched in KIRP, consistent with renal tubular lineage identity (64).

To further examine whether these cancer-specific TF-centered subnetworks reflected lineage identities, we mapped each TF hub to Human Protein Atlas RNA tissue specificity classifications (**Figure S8**, **Table S8**). LGG-specific TF subnetworks were predominantly enriched for brain-lineage TFs. Across other cancer types, TFs with cancer-specific subnetworks generally reflected several biological aspects: (1) tissue-lineage TFs associated with cell-of-origin identity; (2) cancer-testis antigens, which are normally testis-restricted but can become epigenetically reactivated in tumors (65); (3) neural-enriched TF genes potentially reflecting cancer-promoting roles of neural stemness and developmental programs in non-neural cancers (66,67); and (4) immune or lymphoid TFs, likely associated with tumor microenvironment composition. The co-occurrence of these distinct biological programs of cancer-specific TF subnetworks supports the interpretation that the G2G network captures transcriptional coordination beyond cell-of-origin identity, including oncofetal reprogramming and cancer stemness.

Taken together, these results show that methylation-adjusted G2G networks expose biologically interpretable, cancer-specific transcriptional regulatory programs. These networks recapitulate known lineage-associated and cancer-associated TF programs, while also providing a systematic basis for prioritizing cancer-specific TFs and characterizing their co-regulatory architectures.

## Discussion

In this work, we demonstrate that ultrahigh-dimensional multi-omic partial correlation network estimation can be made practical through the example of pan-cancer transcriptome-methylome data. By exploiting the row-separable ACCORD pseudolikelihood formulation, *P*_2_ stabilization, semismooth Newton optimization and a PyTorch implementation for CPU/GPU environments, fastACCORD enabled estimation of joint networks, each input data containing more than 300,000 molecular features, in TCGA cohorts spanning 16 cancer types. This computational capability allowed us to analyze methylation-expression dependencies and methylation-adjusted G2G architectures within the same high-dimensional residual association structure, rather than relying on pairwise methylation-expression associations or post hoc annotation of mRNA-only networks.

From a computational perspective, fastACCORD addresses a key limitation of conventional (e.g. Gaussian) likelihood-based sparse precision matrix workflows: their dependence on matrix inversion operations over the full feature space. In contrast, the ACCORD pseudolikelihood requires inversion of the diagonal component of the reparameterized precision structure only and decomposes naturally across rows. This separability allows row blocks to be processed independently on commodity CPU/GPU hardware. The additional *l*_2_ regularization was particularly useful for multi-omic data, where locally correlated CpG methylation features and correlated expression programs can produce strong collinearity. In the TCGA application, the local correlation structure of CpG methylation features likely contributed to sparse and structured network estimates, although fastACCORD itself is not limited to DNA methylation data.

The joint networks first revealed a pan-cancer landscape of methylation-expression dependencies. The sign of cis-M2G associations depended strongly on genomic location as expected from the literature: promoter proximal methylation was predominantly negatively associated with mRNA expression, consistent with promoter methylation-associated repression, whereas gene-body methylation showed both positive and negative associations. The cross-cancer variation in M2G connectivity suggests that tissue- or cancer-specific epigenetic dysregulation contributes to methylation-expression coupling beyond differences in cohort size. Recurrent negative promoter proximal M2G associations were enriched for metabolic genes, including amino acid catabolism, fatty acid and lipid metabolism, and carbohydrate metabolism. Genes such as THNSL2 and LDHB illustrate how recurrent methylation-associated repression may be linked to tumor metabolic remodeling.

The G2G component of the joint model showed that DNA methylation covariation substantially reshapes gene co-expression architecture. Across all cancers, methylation-adjusted G2G networks were sparser than networks without the adjustment, suggesting that many gene-to-gene associations are no longer retained once measured DNA methylation features are included in the partial correlation modelling. The magnitude of this reduction varied across cancers, reflecting a combination of sample size, cancer-specific methylation structure and molecular heterogeneity. Cross-cancer comparisons further suggested that methylation adjustment can reveal shared gene co-expression architecture that may be masked by cancer-specific methylation patterns, as seen in the increased similarity between LUAD and LUSC after adjustment. Hub genes in methylation-adjusted networks were also enriched for CRISPR-essential genes, suggesting that these networks better capture functionally important hubs linked to tumor-cell survival than mRNA-only networks.

The interpretation of edges altered by methylation adjustment depends on the underlying causal architecture. If DNA methylation acts as a confounder of two genes, including methylation features can remove a non-transcriptional source of co-expression. If methylation lies on a causal path from a regulator TF to its target gene, adjustment may instead attenuate a real, methylation-mediated regulatory signal. Other methylation-linked processes, including subtype structure, tumor purity and cell-composition differences, may also contribute. We therefore do not assign a causal role to individual removed or retained edges. Instead, we interpret methylation-adjusted G2G networks as residual co-expression structures not explained by measured DNA methylation covariation, and we evaluated their aggregate properties using orthogonal regulatory and functional evidence.

Consistent with this view, methylation-adjusted G2G networks showed stronger enrichment for TF-associated regulatory evidence than mRNA-only networks. Adjusted networks contained a higher proportion of ReMap-supported TF-target edges and gene pairs supported by curated co-regulon annotations, and retained edges showed stronger partial correlations than edges removed by methylation adjustment. These observations suggest that accounting for DNA methylation covariation sharpens the TF-associated component of G2G co-expression networks. Importantly, these annotations do not convert an undirected partial correlation network into a causal GRN. Rather, they show that the retained gene-gene associations are more concentrated among gene pairs supported by independent TF-binding and regulon evidence.

TF-centered subnetworks in the methylation-adjusted G2G networks revealed highly cancer-specific co-expression architectures. Many cancer-specific TF subnetworks recapitulated known lineage-associated and cancer-associated TF programs, including ESR1 and FOXA1 in BRCA, AR in PRAD, HMG/Sox family TFs in LGG and renal-lineage-associated homeodomain TFs in KIRP. Beyond classical lineage regulators, cancer-specific TF subnetworks also included cancer-testis antigens, neural developmental TFs, and immune-lineage TFs, the latter likely reflecting tumor-infiltrating immune cell signals in bulk RNA-seq. Notably, cancer-testis antigen and neural-lineage TFs are normally silenced or tissue-restricted outside their lineage of origin. Their formation of cancer-specific TF co-expression hubs therefore suggests that subnetwork structure captures oncofetal and developmental transcriptional programs that are absent or restricted in normal tissues. For example, SALL4, a testis-enriched TF silenced in normal adult tissue, formed a LIHC-specific subnetwork consistent with its established oncofetal role in aggressive HCC (68), with its subnetwork recapitulating classical HCC biomarkers (AFP, GPC3) and an IGF2/IGF2BP1/IGF2BP2 oncofetal signature (**Figure S9**). Collectively, TF co-expression subnetworks reflect tissue identity, lineage reprogramming, and oncofetal reactivation in a cancer type-specific manner.

More broadly, the networks reported here should be interpreted as partial correlation networks: sparse maps of residual linear associations among molecular variables. They are not complete causal regulatory graphs. Rather, they are expected to contain a broad superset of relationships, including but not limited to direct TF-target relationships. For this reason, fastACCORD is best used as a scalable graph discovery and prioritization framework. A proper GRN reconstruction should incorporate complementary information such as regulatory sequence features, ChIP-seq evidence, and tissue- or cell-type-specific expression levels.

Despite the advantages of fastACCORD and its utility demonstrated in this work, we acknowledge that the approach has a few practical constraints. First, fastACCORD currently operates on continuously scaled variables and does not directly model binary, categorical or ordinal features. Extending the framework to mixed graphical models will be important for incorporating mutation status, copy-number events, clinical annotations and other discrete variables. Second, although fastACCORD reduces computational barriers, practical performance still depends on feature dimension, selected graph density, row-block size, active-set size and available CPU/GPU memory. Users may need to tune implementation parameters for their specific data and hardware environment.

There are several biological and annotation-related limitations as well. First, TCGA profiles are derived from bulk tumors, so inferred networks can be a consequence of multiple factors such as tumor-intrinsic regulation, tumor purity, immune or stromal composition, and molecular subtype structure. Second, TF binding resources (e.g. ReMap) and motif resources (e.g. JASPAR) are biased toward well-studied TFs and well-profiled cell types, limiting uniform validation across the full TF repertoire. Third, TCGA cohort sizes vary substantially across cancer types, and smaller cohorts have reduced power to support stable edges in high-dimensional partial correlation networks. Differences in cohort size and graph density may therefore affect cross-cancer comparisons of network size, similarity and TF subnetwork specificity. Fourth, our TF-centered analyses are based primarily on mRNA-level expression similarity. As mentioned earlier, post-transcriptionally regulated TFs, primarily activated through protein stabilization rather than transcriptional change, may therefore be underrepresented in our co-expression-based subnetworks. Fifth, adjusted and unadjusted G2G networks differ in their conditioning sets and feature dimensions, which can affect graph density and edge selection. We therefore interpret their differences at the aggregate network level rather than as definitive edge-level causal classifications. Finally, although miRNA expression was included in the full integrated network, the present analyses focused on M2G and G2G subnetworks. Future work could examine miRNA-centered regulatory subnetworks.

For future development, several extensions would broaden the utility of fastACCORD-based network analysis. One direction is to integrate additional molecular layers, including chromatin accessibility, histone marks, genetic variation, copy-number alterations and proteomic measurements. Another is to develop weighted or block-specific regularization schemes so that inter-modality edges are not overly penalized relative to dense within-modality correlation structure. A third direction is to combine ultrahigh-dimensional partial correlation networks with causal discovery, perturbation data or prior regulatory knowledge to orient selected edges or infer partially directed graph structures. These developments would move multi-omic network analysis beyond undirected residual association maps while preserving the scalability needed for genome-scale molecular data.

In summary, fastACCORD provides a scalable framework for ultrahigh-dimensional partial correlation modeling and multi-omic data integration. Applied to TCGA transcriptome-methylome data, fastACCORD produced a pan-cancer network resource that captures methylation-expression dependencies, methylation-adjusted gene-gene co-expression and cancer-specific TF-associated co-regulatory landscapes. This resource offers a practical foundation for prioritizing cancer-specific TFs, exploring epigenomically modulated co-expression programs and designing follow-up regulatory analyses in cancer.

## Data availability

fastACCORD is available at https://github.com/accord-dev/fastACCORD. A user manual for the program is also available in this repository. G2G network visualizer is available at https://github.com/SLINGhub/ACCORD-TCGA-G2G_network_visualizer. Data containing the full network (with M2G edges), adjusted G2G networks, and unadjusted G2G networks for all cancer types are deposited in Zenodo (10.5281/zenodo.22183622).

## Author contribution

SL, DK, SO and JW contributed to algorithm development. SL designed and implemented the fastACCORD software. QZ curated data, performed pan-cancer network analysis, and developed network visualizer application. SL and DK conducted simulation studies for performance benchmarking. SL, QZ and HC drafted manuscript and all authors contributed to manuscript writing. SO, JW and HC supervised the project.

## Acknowledgment

The results published here are in whole or part based upon data generated by the TCGA Research Network: https://www.cancer.gov/tcga. All data used in this work was downloaded through NCI Genomic Data Commons (69). We used Claude for the development of network visualizer, ChatGPT 5-6 in improving the readability and language of the manuscript, and FigureLabs for creating the graphical abstract. After the use of these tools, all authors reviewed and edited the content and take full responsibility for the content of the publication.

## Abbreviations

ACCORD: Asymmetric convex partial correlation selection method
BLCA: Bladder urothelial carcinoma
BRCA: Breast invasive carcinoma
CESC: Cervical squamous cell carcinoma and endocervical adenocarcinoma
DAG: Directed acyclic graph
G2G: Gene-to-gene
GRN: Gene regulatory network
HNSC: Head and neck squamous cell carcinoma
HPC: High-performance computing
KIRC: Kidney renal clear cell carcinoma
KIRP: Kidney renal papillary cell carcinoma
LGG: Brain lower grade glioma
LIHC: Liver hepatocellular carcinoma
LUAD: Lung adenocarcinoma
LUSC: Lung squamous cell carcinoma
M2G: Methylation-to-gene
MPI: Message-passing interface
PRAD: Prostate adenocarcinoma
SARC: Sarcoma
STAD: Stomach adenocarcinoma
TCGA: The Cancer Genome Atlas
TF: Transcription factor
TGCT: Testicular germ cell tumors
THCA: Thyroid carcinoma
TSS: Transcription start site
UCEC: Uterine corpus endometrial carcinoma
VRAM: Video random access memory

## Supplementary Data

Supplementary Data are available online. Supplementary materials include additional methodological details and computational results, Supplementary Figures S1-S9, and Supplementary Tables S1-S8.

