## Supplementary Information for "fastACCORD enables ultrahigh-dimensional partial correlation modeling for multi-omic data integration"

#### Optimization of fastACCORD with the semismooth Newton method

The objective of fastACCORD is to find a $p \times p$ matrix $\boldsymbol{\Omega}$ that minimizes

$$-\log \det \boldsymbol{\Omega}_{\mathbf{D}} + \frac{1}{2}\mathrm{tr}(\boldsymbol{\Omega}^{T}\boldsymbol{\Omega}\mathbf{S}) + \lambda_1\|\boldsymbol{\Omega}_{-D}\|_1 + \frac{\lambda_2}{2}\|\boldsymbol{\Omega}\|_2^2,$$

where $\mathbf{S} = \mathrm{n}^{-1}\mathbf{X}^{\mathbf{T}}\mathbf{X}$ is the sample covariance, and $\boldsymbol{\Omega}_{\mathbf{D}}$ and $\boldsymbol{\Omega}_{-D}$ denotes a matrix only with and without diagonal parts of $\boldsymbol{\Omega}$, respectively. Note that this objective function can be decomposed into

$$\sum_{i=1}^{p}\left(-\log\det\boldsymbol{\Omega}_{i,i} + \frac{1}{2}\mathrm{tr}\left(\boldsymbol{\Omega}_i^T\boldsymbol{\Omega}_i\mathbf{S}\right) + \lambda_1\left\|\boldsymbol{\Omega}_{i,-i}\right\|_1 + \frac{\lambda_2}{2}\|\boldsymbol{\Omega}_i\|_2^2\right),$$

where $\boldsymbol{\Omega}_i$ denotes the $i$-th row of $\boldsymbol{\Omega}$, $\boldsymbol{\Omega}_{i,i}$ and $\boldsymbol{\Omega}_{i,-i}$ denotes the diagonal and the off-diagonal parts of $\boldsymbol{\Omega}_i$, respectively. Without loss of generality, we will first focus on optimizing the objective above with respect to $i = 1$ employing the semismooth Newton method, a second-order approach for convex problems with non-smooth yet semi-smooth objective functions [1,2]. Let $\omega = (\omega_1, \omega_{-1}^T)^T$, where $\omega_1 = \boldsymbol{\Omega}_{1,1}$ and $\omega_{-1} = \boldsymbol{\Omega}_{1,-1}$. Also, let

$$g(\omega) = \tfrac{1}{2}\omega^T\mathbf{S}\omega + \tfrac{\lambda_2}{2}\|\omega\|_2^2, \quad h(\omega) = -\log\omega_1 + \lambda_1\|\omega_{-1}\|_1.$$

The Karush-Kuhn-Tucker optimality condition for $g(\omega) + h(g)$ can be expressed as

$$(\mathbf{S} + \lambda_2\mathbf{I})\omega^* + d^* = 0, \qquad d^* \in \partial h(\omega^*)$$

Where $\partial h$ denotes the subdifferential $h$. This is equivalent to the root of the map $F: \mathbb{R}^{2p} \to \mathbb{R}^{2p}$ defined as,

$$F(\omega, \mathrm{d}) = \begin{pmatrix} (\mathbf{S} + \lambda_2\mathbf{I})\omega + \mathrm{d} \\ \omega_1 - \mathbf{prox}_{-\log(\cdot)}(\omega_1 + \mathrm{d}_1) \\ \omega_{-1} - \mathrm{T}_\lambda(\omega_{-1} + \mathrm{d}_{-1}) \end{pmatrix}, \quad \mathbf{prox}_{-\log(\cdot)}(\mathrm{x}) = \tfrac{1}{2}\left(\mathrm{x} + \sqrt{\mathrm{x}^2 + 4}\right)$$

Where $T_\lambda(\omega_{-1}) = \left(\mathrm{sign}(\omega_j)\max(|\omega_j| - \lambda, 0)\right)_{j\neq 1}$ is an element-wise soft-thresholding operator. Denoting $x = (\omega, d)$, the Newton iterate for finding the root of $F$ is given as

$$x^{k+1} = x^k - G(x^k)^{-1}F(x^k)$$

where $G$ is the nonsingular generalized Jacobian matrix of $F$. One possible formulation of $G$ is

$$G = \begin{pmatrix} \mathbf{S} + \lambda_2\mathbf{I} & \mathbf{I} \\ \mathbf{I} - \mathbf{J} & -\mathbf{J} \end{pmatrix},$$

where $\mathbf{J} = \mathrm{diag}(\mathbf{J}_{ii})$ is a diagonal matrix with

$$\mathbf{J}_{11} = \frac{1}{2} + \frac{\omega_1 + d_1}{\sqrt{(\omega_1 + d_1)^2 + 4}}$$

and

$$\mathbf{J}_{\mathrm{ii}} = \begin{cases} 1, & |\omega_i + d_i| > \lambda_1 \\ 0, & |\omega_i + d_i| \leq \lambda_1 \end{cases}$$

for $i = 2, \cdots, p$. Next, define the active and inactive set for the $k$-th iterate $(\omega^k, d^k)$ as

$$\mathbf{A}^{\mathrm{k+1}} \coloneqq \{i \in \{2, \dots, p\}: |\omega_i^k + d_i^k| > \lambda_1\},\ \mathbf{I}^{\mathrm{k+1}} \coloneqq \{i \in \{2, \dots, p\}: |\omega_i^k + d_i^k| \leq \lambda_1\}.$$

We also define $\overline{\boldsymbol{A}}^{\mathrm{k+1}} = \{1\} \cup \mathbf{A}^{\mathrm{k+1}}$ for convenience. Provided that $G$ is invertible, the equation $G(x^k)x^{k+1} = G(x^k)x^k - F(x^k)$ yields

$$\omega_{\mathbf{I}^{\mathrm{k+1}}}^{k+1} = 0, \qquad d_{\mathbf{A}^{\mathrm{k+1}}}^{k+1} = \begin{cases} \lambda_1 & if\ \omega_i + d_i > \lambda_1, \\ -\lambda_1 & if\ \omega_i + d_i < -\lambda_1, \end{cases}$$

$$\left[\begin{pmatrix} \mathbf{S}_{11} & \mathbf{S}_{1\mathbf{A}^{\mathrm{k+1}}} \\ \mathbf{S}_{\mathbf{A}^{\mathrm{k+1}}1} & \mathbf{S}_{\mathbf{A}^{\mathrm{k+1}}\mathbf{A}^{\mathrm{k+1}}} \end{pmatrix} + \lambda_2 \mathbf{I}\right] \begin{pmatrix} \omega_1^{k+1} \\ \omega_{\mathbf{A}^{\mathrm{k+1}}}^{k+1} \end{pmatrix} + \begin{pmatrix} d_1^{k+1} \\ d_{\mathbf{A}^{\mathrm{k+1}}}^{k+1} \end{pmatrix} = 0,$$

$$\left(z^k + \frac{1}{2}\right)\omega_1^{k+1} + \left(z^k - \frac{1}{2}\right)d_1^{k+1} = z^k(\omega_1^k + d_1^k) - \frac{\sqrt{\left(\omega_1^k + d_1^k\right)^2 + 4}}{2},$$

where $z^k = -\frac{\omega_1^k + d_1^k}{2\sqrt{(\omega_1^k + d_1^k)^2 + 4}}$. Substituting $d_1^{k+1}$ with the last equation, the semismooth Newton iterate reduces to solving the linear system

$$\left[\begin{pmatrix} \mathbf{S}_{11} + \mathrm{b}^{\mathrm{k}} & \mathbf{S}_{1\mathbf{A}^{\mathrm{k+1}}} \\ \mathbf{S}_{\mathbf{A}^{\mathrm{k+1}}1} & \mathbf{S}_{\mathbf{A}^{\mathrm{k+1}}\mathbf{A}^{\mathrm{k+1}}} \end{pmatrix} + \lambda_2 \mathbf{I}\right] \begin{pmatrix} \omega_1^{k+1} \\ \omega_{\mathbf{A}^{\mathrm{k+1}}}^{k+1} \end{pmatrix} = \begin{pmatrix} c^k \\ \pm\lambda_1 \end{pmatrix},$$

$$\mathrm{b}^{\mathrm{k}} = -\left(z^k - \frac{1}{2}\right)^{-1}\left(z^k + \frac{1}{2}\right), c^k = -\left(z^k - \frac{1}{2}\right)^{-1}\left(z^k(\omega_1^k + d_1^k) - \frac{\sqrt{\left(\omega_1^k + d_1^k\right)^2 + 4}}{2}\right).$$

Note that $b^k$ and $c^k$ are explicitly defined with $(\omega_1^k, d_1^k)$, and $b^k$ is always positive.

There are several remarks for this semismooth Newton iteration. First, the invertibility of the generalized Jacobian $G$ is equivalent with the non-singularity of the linear system to solve. For the original ACCORD with $\lambda_2 = 0$, the matrix $\mathbf{S}_{\overline{\boldsymbol{A}}^{\mathrm{k+1}}} = (1/n)\mathbf{X}_{\overline{\boldsymbol{A}}^{\mathrm{k+1}}}^{\mathbf{T}}\mathbf{X}_{\overline{\boldsymbol{A}}^{\mathrm{k+1}}}$ is singular with a non-trivial null space whenever size of the active set $|\overline{\boldsymbol{A}}^{\mathrm{k+1}}|$ exceeds the sample size $n$, possibly making the update intractable. However, the coefficient matrix of the linear system is always positive definite for $\lambda_2 > 0$. Hence, imposing the positive Frobenius norm guarantees the successful computation of the semismooth Newton update regardless of the current active set size. Nevertheless, an excessively large size of $|\mathbf{A}^{\mathrm{k+1}}|$ can pose a significant computational bottleneck in practice even for $\lambda_2 > 0$, since solving the linear system above demands time complexity of $O(|\mathbf{A}^{\mathrm{k+1}}|^3)$ and space complexity of $O(|\mathbf{A}^{\mathrm{k+1}}|^2)$. At a scale of $p$ where operations on the full $p \times p$ matrix become computationally intractable, the iteration likewise becomes intractable as $|\mathbf{A}^{\mathrm{k+1}}|$ approaches $p$. In such case, we can still adopt the proximal gradient descent method in [3] converging to the optimal solution in a linear rate, which is

$$\omega^{k+1} = \mathbf{prox}_{\tau \mathrm{h}}(\omega^k - \tau \nabla g(\omega^k)),$$

where $\nabla g(\omega) = \mathbf{S}\omega$ and $\mathbf{prox}_{\mathrm{h}}$ is a soft-thresholding operator for off-diagonal entries and $\mathbf{prox}_{-\mathbf{log}(\cdot)}$ for diagonal entries. If the optimal solution is sparse with nonzero entries less than $n$, we can expect that the proximal gradient steps will eventually approach to a region where the semismooth Newton update is also feasible. Finally, although the semismooth Newton method enjoys a local quadratic convergence rate, it does not necessarily guarantee the global convergence—starting from an ill-conditioned initial point can blow up the iterate. To ensure the stable convergence of the algorithm, we have also adopted the damped semismooth Newton iteration incorporating a line search, which is fully integrated into our software implementation. Refer to [1] for details.

### Computational workflow and results

We describe the workflow for predicting networks from 16 TCGA data sets, encompassing both DNA methylation-adjusted and unadjusted networks. Prior to the analysis, DNA methylation and mRNA and microRNA expression values were preprocessed and standardized to construct a $n \times p$ NumPy array for each type of cancer. Using our fastACCORD software implementation, the ACCORD estimates were computed across a predefined grid of distinct $\lambda_1$ values. The estimate from the previous grid point was used as a warm-start initial value if available, and the identity matrix was used otherwise.

To facilitate distributed processing in a multi-GPU environment, we split the estimation of the full $p \times p$ matrix parameter into 500 to 1,000 row blocks, depending on the size of the estimate. Each resulting task was then allocated to and executed on a single GPU across all available GPUs in the environment. For the unadjusted network only with mRNA features, we split the task in 4 row blocks. The table below reports the **average time per GPU** in completing all the allocated tasks on computing the selected estimate for each TCGA data set. For the hardware and software stack, we employed four NVIDIA TITAN V GPU alongside PyTorch v2.10.0 and CUDA v13.0.

| | | Number of features | | Average time per GPU (s) | |
|---|---|---|---|---|---|
| Data | n | Total | mRNA | Adjusted | Unadjusted |
| BLCA | 398 | 349,877 | 15,935 | 1,796 | 55 |
| BRCA | 599 | 317,771 | 16,325 | 1,995 | 69 |

| | | | | | |
|---|---|---|---|---|---|
| CESC | 303 | 326,718 | 16,078 | 510 | 45 |
| HNSC | 515 | 348,721 | 16,167 | 687 | 66 |
| KIRC | 313 | 347,160 | 16,409 | 584 | 71 |
| KIRP | 274 | 372,387 | 16,017 | 667 | 65 |
| LGG | 512 | 347,095 | 16,611 | 660 | 86 |
| LIHC | 367 | 325,322 | 15,341 | 580 | 63 |
| LUAD | 443 | 350,002 | 16,363 | 645 | 62 |
| LUSC | 366 | 331,278 | 16,590 | 1,359 | 50 |
| PRAD | 489 | 360,111 | 16,467 | 2,423 | 84 |
| SARC | 257 | 325,943 | 15,706 | 937 | 42 |
| STAD | 363 | 327,766 | 16,841 | 1,920 | 58 |
| TGCT | 150 | 388,251 | 16,983 | 716 | 71 |
| THCA | 504 | 347,318 | 16,183 | 752 | 88 |
| UCEC | 425 | 335,937 | 15,953 | 1,910 | 56 |

#### Supplementary Figures

**Figure S1**. Partial correlations against genomic distances between connected nodes. LUAD is shown as a representative example.

**Figure S2**. KEGG pathway enrichment of genes with negative cis-M2G associations at promoter and gene body regions across cancers (Full version of Figure 2C)

**Figure S3**. KEGG pathway enrichment of genes with positive cis-M2G associations at the gene body across cancers.

**Figure S4**. Gene-level Jaccard similarity between LUSC and LUAD co-expression networks before and after methylation adjustment.

**Figure S5**. Distribution of node degrees for all genes (gray) and CRISPR-essential genes (Chronos score < −1, blue) within the networks.

**Figure S6**. Cancer type-specific ChIP-seq-validated TF-target annotation rate in methylation-adjusted and unadjusted co-expression networks. For each cancer type, only ChIP-seq experiments from selected cell lines were included (Table S4). The number of unique TFs profiled in each cancer-specific background is shown in parentheses.

**Figure S7**. Proportion of co-expression edges supported by ≥3 shared upstream TFs in methylation-adjusted and unadjusted networks.

**Figure S8**. Lineage classification of 653 cancer-specific TF subnetwork hubs. Details in Table S8. Confirmed lineage: TF is HPA tissue-specific for the exact tissue of origin of the peak cancer. Adjacent/related lineage: TF specific to an anatomically adjacent, developmentally related, or reproductive tissue. Cancer-testis antigen: Testis-restricted TF epigenetically de-repressed in cancer. Neural reprogramming: Brain-specific TF active in cancers with neural-crest origin or neural-like plasticity. Immune/TME: Bone marrow/lymphoid TF reflecting immune cell infiltration in the tumor microenvironment. Unclear: TF enriched in broadly shared or unrelated tissues; No HPA tissue-specific classification: TF not classified as tissue-specific/enhanced by HPA.

**Figure S9**. Subnetwork for SALL4 in LIHC. The network was generated using the G2G network visualizer developed in this paper.

#### Supplementary Tables

**Table S1.** TCGA data resources and gene co-expression network metrics

**Table S2.** Pan-cancer enrichment of recurrent negative cis DNA methylation-to-mRNA associations at gene promoters (present in ≥3 cancer types)

**Table S3.** Cancer type-matched DepMap cell lines selected for CRISPR gene dependency analysis

**Table S4.** Cancer type-matched cell lines and tissues selected for ReMap ChIP-seq background construction

**Table S5.** Most extensively ChIP-seq profiled TFs in cancer type-matched cell lines from ReMap

**Table S6.** Cancer-specific TF co-expression hub metrics across 16 TCGA cancer types

**Table S7.** Curated TF-cancer associations from established cancer drivers and master regulators

**Table S8**. Lineage specificity for TFs with cancer-specific subnetworks
