## Supplementary Figures 1-9 for "fastACCORD enables ultrahigh-dimensional partial correlation modeling for multi-omic data integration"

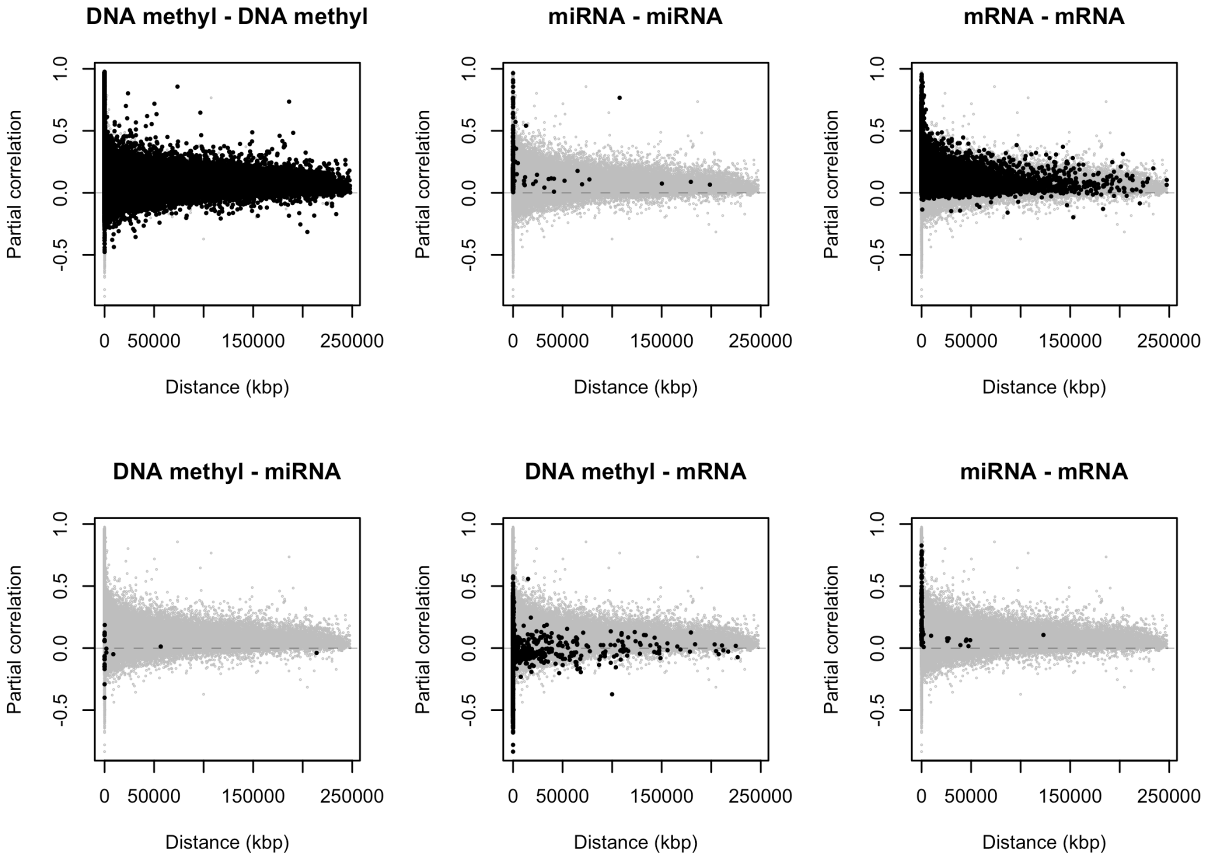


**Figure S1.** Partial correlations against genomic distances between connected nodes. LUAD is shown as a representative example.

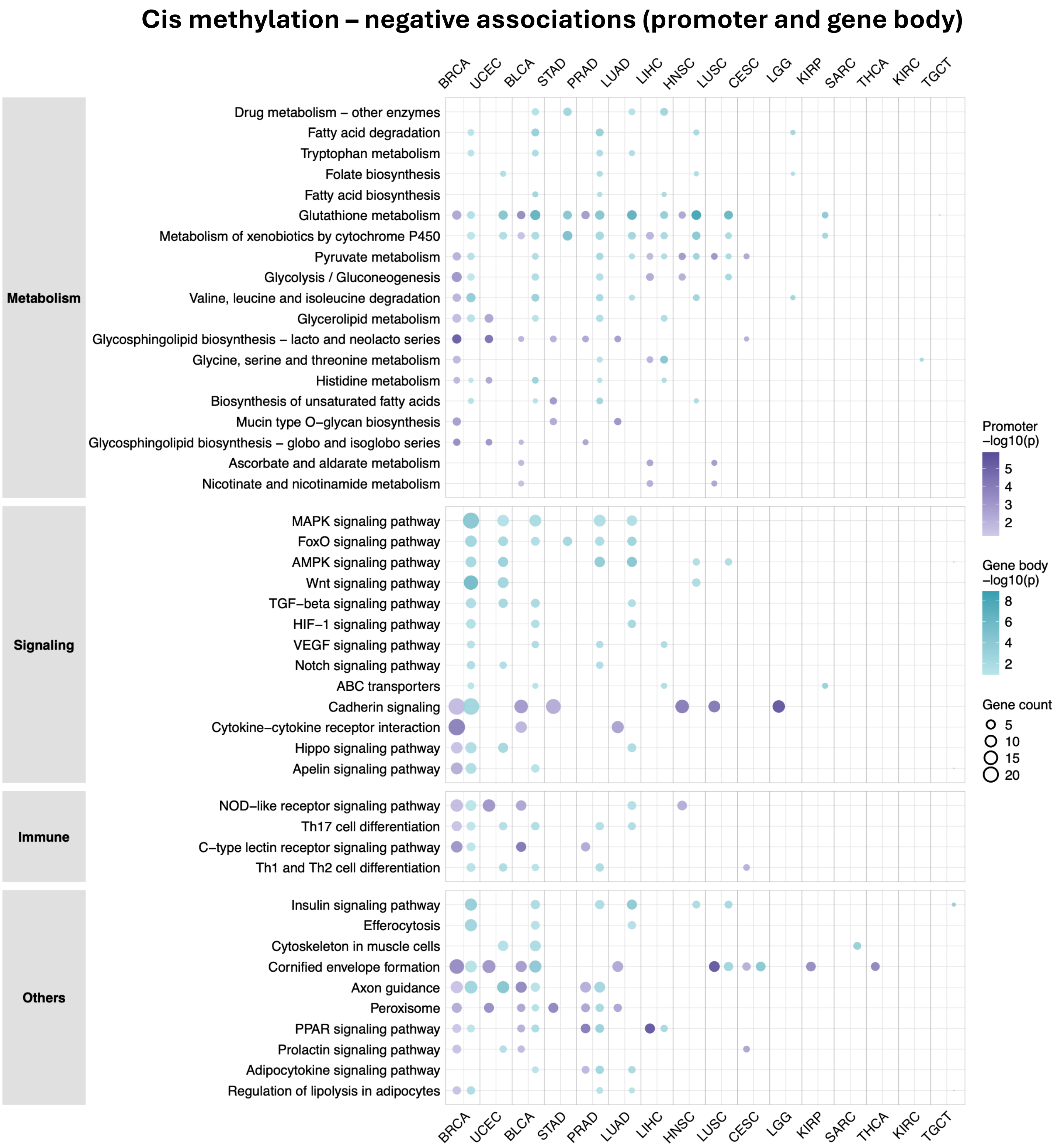


**Figure S2.** KEGG pathway enrichment of genes with negative cis M2G associations at promoter and gene body regions across cancers (Full version of Figure 2C)

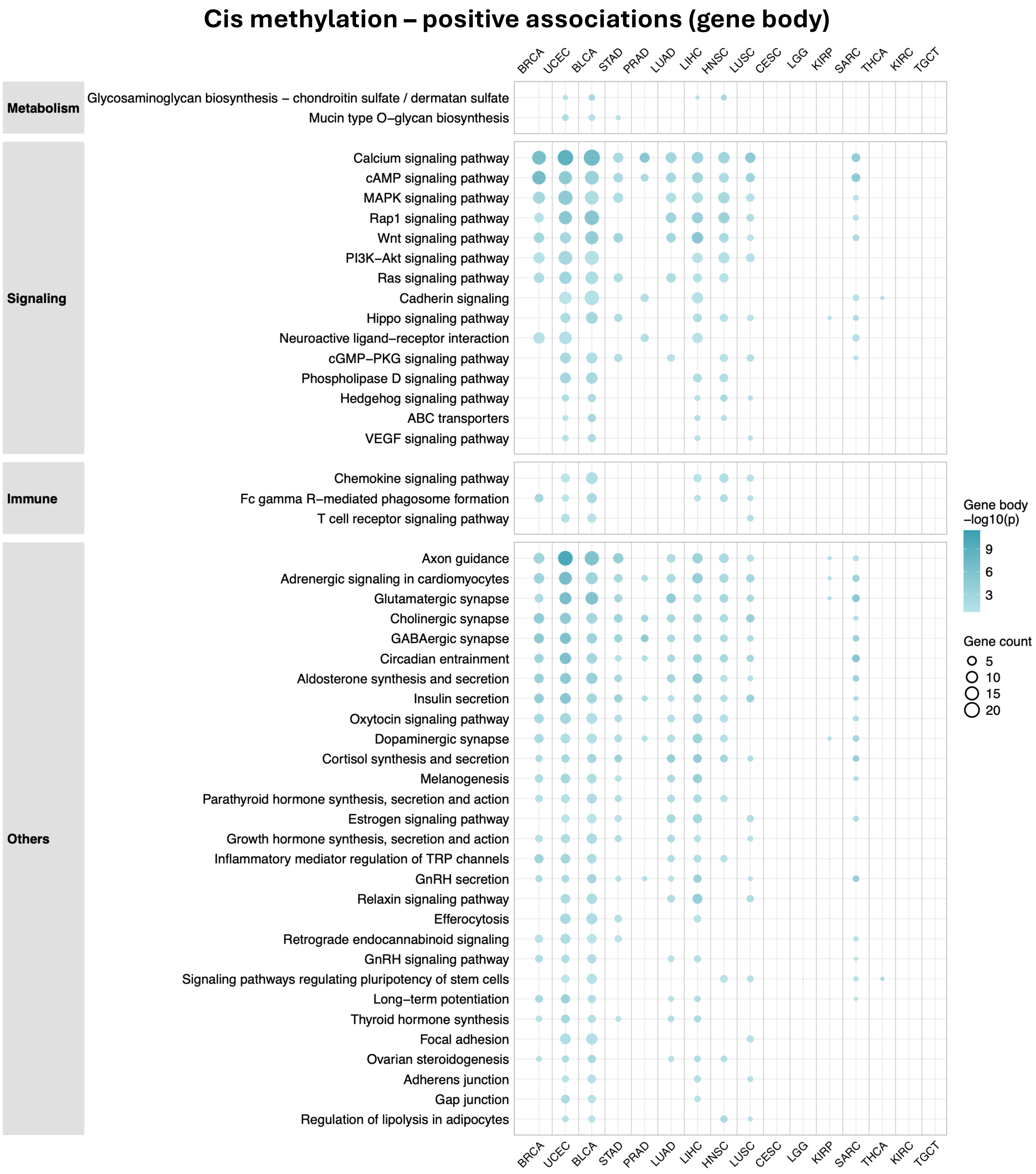


**Figure S3.** KEGG pathway enrichment of genes with positive cis M2G associations at the gene body across cancers.

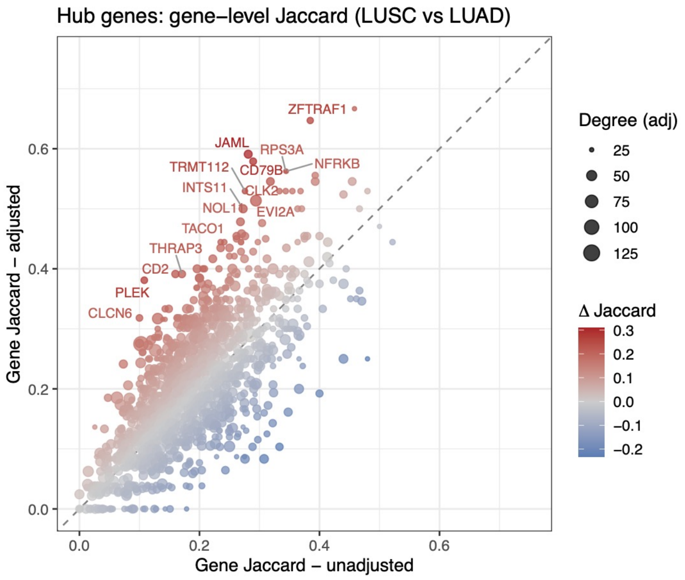


**Top 20 genes**

- Immune cell markers (CD79B, JAML, CD2, PLEK, EVI2A, LST1, ITGAX)
- Mitochondrial/metabolic genes (COX7C, TACO1)
- RNA processing components (TRMT112, LSM5, ESS2)
- Signalling/chromatin regulators (NFRKB, INTS11, DVL3, ZFTRAF1, NOL11, CLCN6, PRR14L)
- Transcription factor (ZNF225)

**Figure S4.** Gene-level Jaccard similarity between LUSC and LUAD co-expression networks before and after methylation adjustment.

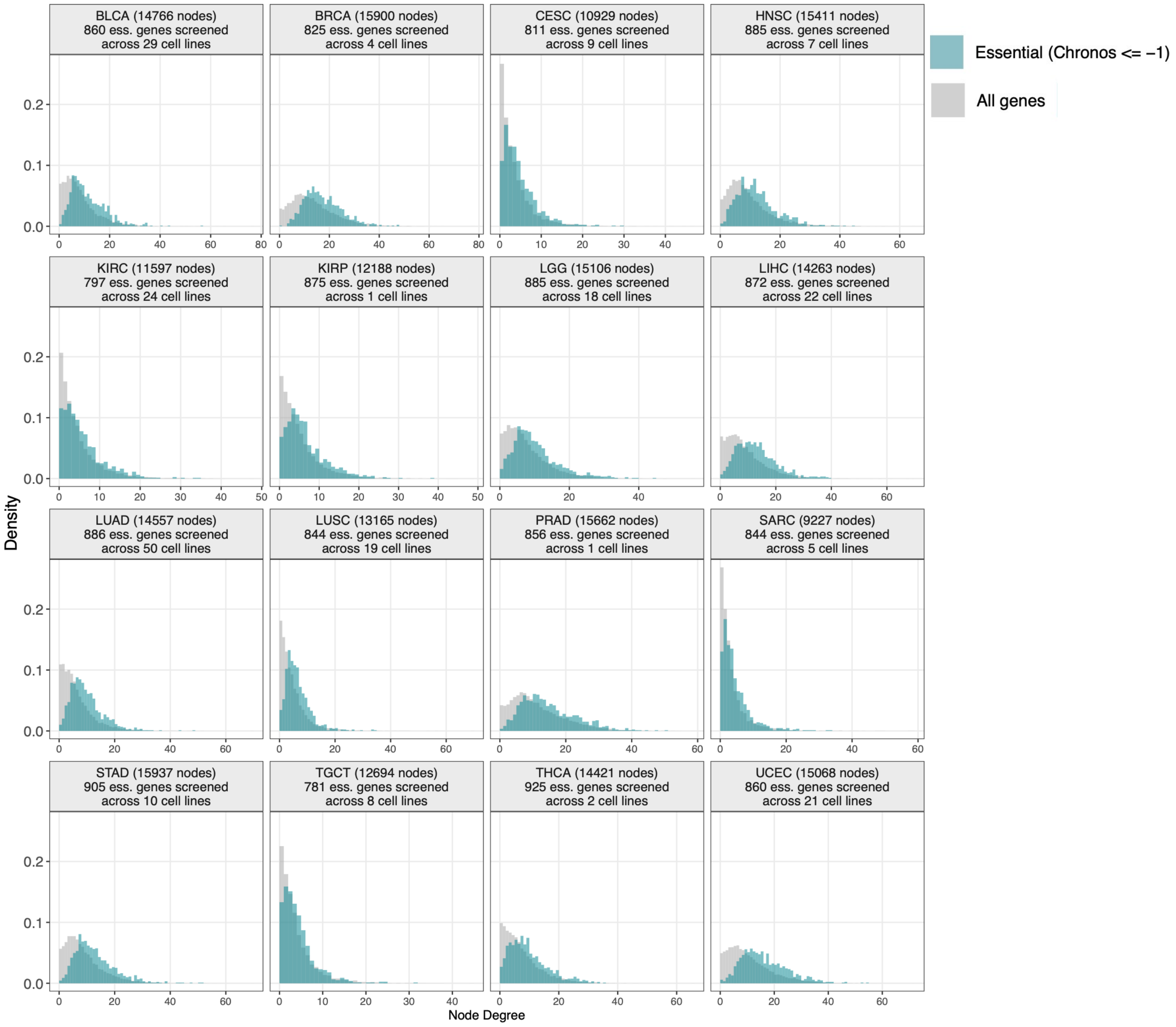


**Figure S5.** Distribution of node degrees for all genes (gray) and CRISPR-essential genes (Chronos score < −1, blue) within the networks.

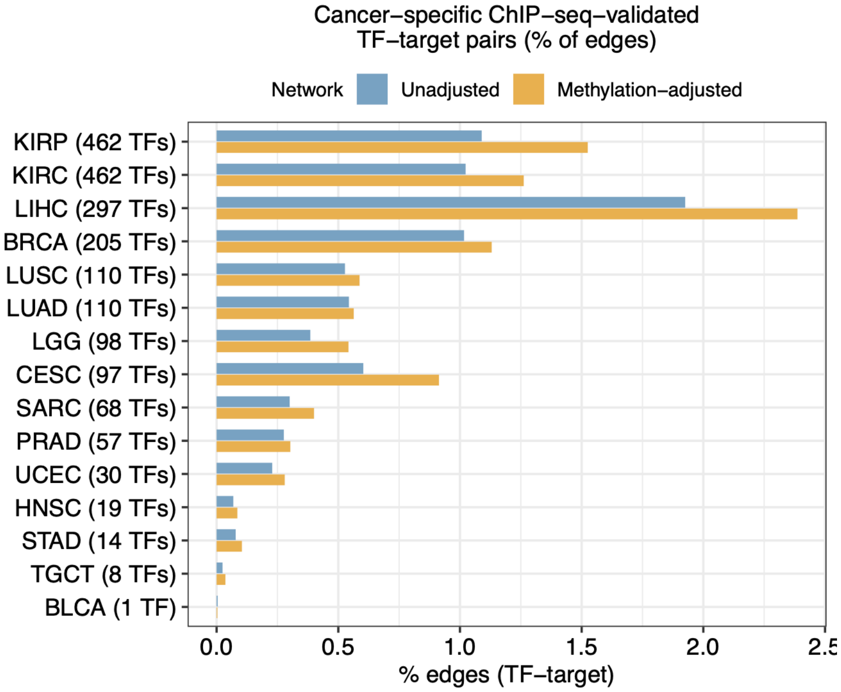


**Figure S6**. Cancer type-specific ChIP-seq-validated TF-target annotation rate in methylation-adjusted and unadjusted co-expression networks. For each cancer type, only ChIP-seq experiments from selected cell lines were included (Table S4). The number of unique TFs profiled in each cancer-specific background is shown in parentheses.

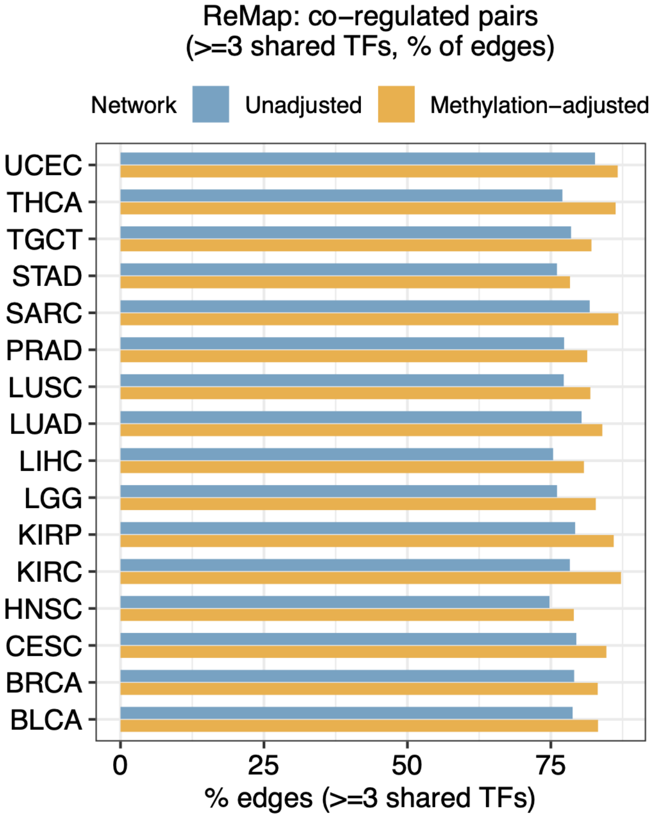


**Figure S7.** Proportion of co-expression edges supported by ≥3 shared upstream TFs in methylation-adjusted and unadjusted networks.

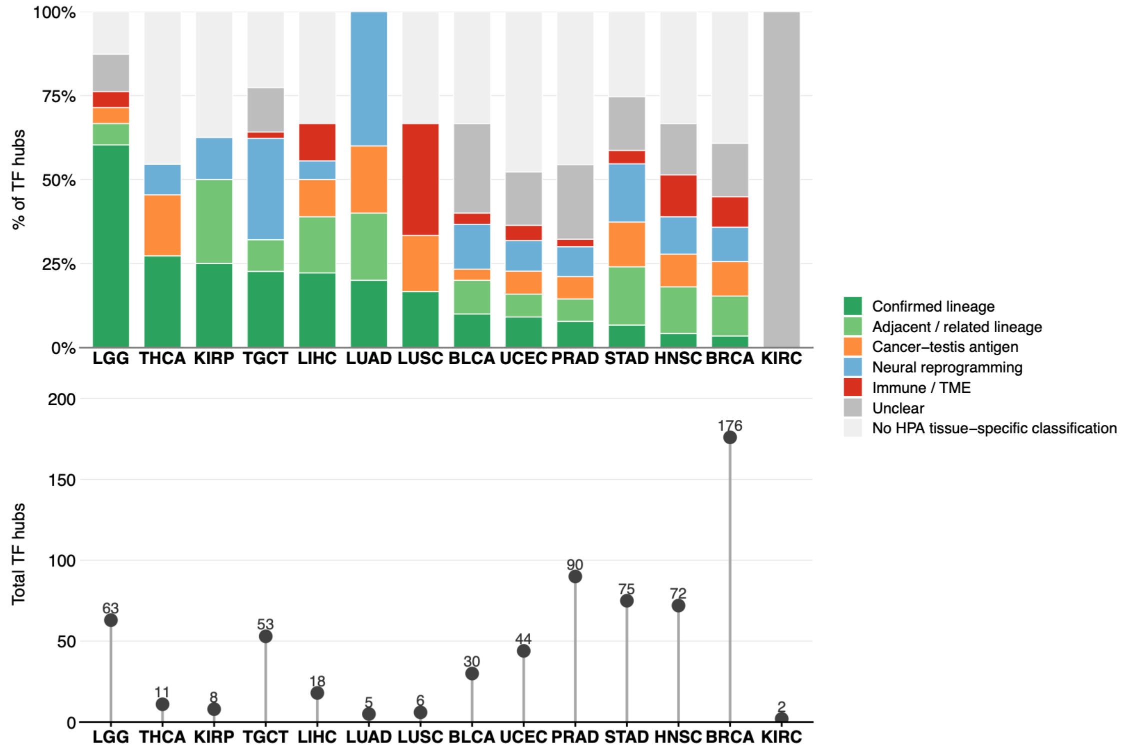


**Figure S8.** Lineage classification of 653 cancer-specific TF subnetwork hubs. Details in Table S8. Confirmed lineage: TF is HPA tissue-specific for the exact tissue of origin of the peak cancer. Adjacent/related lineage: TF specific to an anatomically adjacent, developmentally related, or reproductive tissue. Cancer-testis antigen: Testis-restricted TF epigenetically de-repressed in cancer. Neural reprogramming: Brain-specific TF active in cancers with neural-crest origin or neural-like plasticity. Immune/TME: Bone marrow/lymphoid TF reflecting immune cell infiltration in the tumor microenvironment. Unclear: TF enriched in broadly shared or unrelated tissues; No HPA tissue-specific classification: TF not classified as tissue-specific/enhanced by HPA.

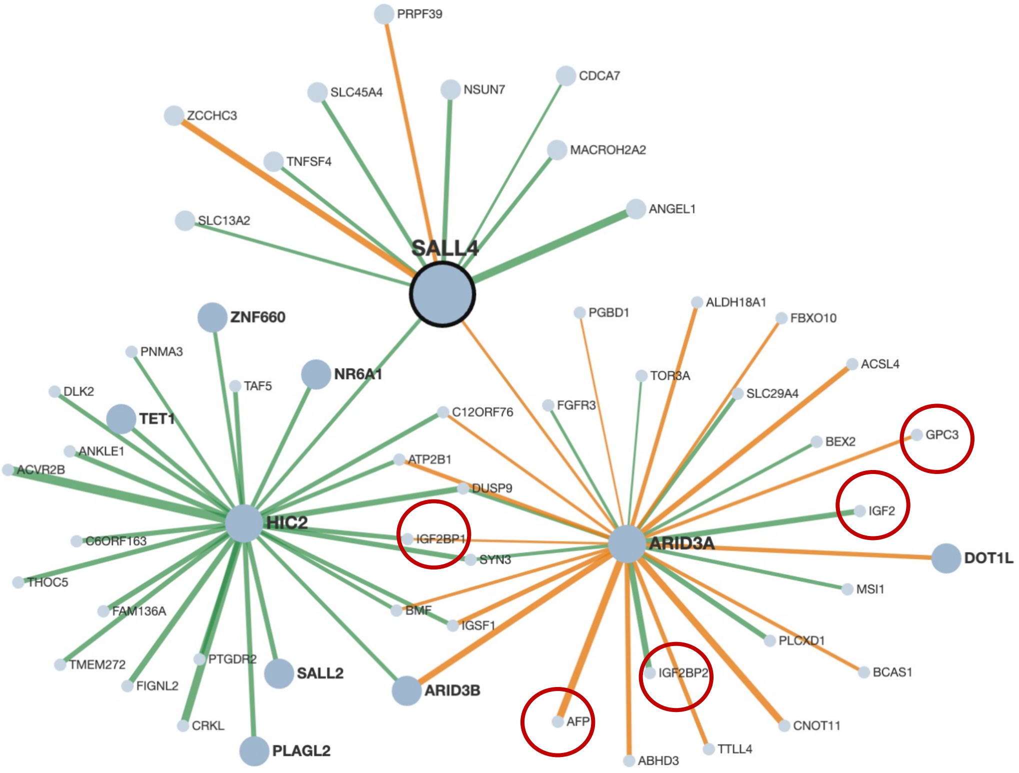


**Figure S9.** Subnetwork for SALL4 in LIHC. The network was generated using the G2G network visualizer developed in this paper.
